# A subset of broadly responsive Type III taste cells contribute to the detection of bitter, sweet and umami stimuli

**DOI:** 10.1101/660589

**Authors:** Debarghya Dutta Banik, Eric D. Benfey, Laura E. Martin, Kristen E. Kay, Gregory C. Loney, Amy R. Nelson, Zachary C. Ahart, Barrett T. Kemp, Bailey R. Kemp, Ann-Marie Torregrossa, Kathryn F. Medler

**Affiliations:** Department of Biological Sciences, University at Buffalo, Buffalo, NY 14260, USA; Department of Psychology, University at Buffalo, Buffalo, NY 14260, USA; Center for Ingestive Behavior Research, University at Buffalo, Buffalo, NY 14260, USA

## Abstract

Taste receptor cells use multiple signaling pathways to detect chemicals in potential food items. These cells are functionally grouped into different types: Type I cells act as support cells and have glial-like properties; Type II cells detect bitter, sweet, and umami taste stimuli; and Type III cells detect sour and salty stimuli. We have identified a new population of taste cells that are broadly tuned to multiple taste stimuli including bitter, sweet, sour and umami. The goal of this study was to characterize these broadly responsive (BR) taste cells. We used an IP3R3-KO mouse (does not release calcium (Ca^2+^) from Type II cells when stimulated with bitter, sweet or umami stimuli) to characterize the BR cells without any potentially confounding input from Type II cells. Using live cell Ca^2+^ imaging in isolated taste cells from the IP_3_R3-KO mouse, we found that BR cells are a subset of Type III cells that respond to sour stimuli but also use a PLCβ3 signaling pathway to respond to bitter, sweet and umami stimuli. Unlike Type II cells, individual BR cells are broadly tuned and respond to multiple stimuli across different taste modalities. Live cell imaging in a PLCβ3-KO mouse confirmed that BR cells use a PLCβ3 signaling pathway to generate Ca^2+^ signals to bitter, sweet and umami stimuli. Analysis of c-Fos activity in the nucleus of the solitary tract (NTS) and short term behavioral assays revealed that BR cells make significant contributions to taste.

---

Chemicals in the oral cavity are detected by taste receptor cells which are grouped together in taste buds found in epithelial specializations called papillae. The current view of taste transduction is that taste buds are comprised of Type I, Type II and Type III taste receptor cells along with basal cells that are precursors for the differentiated cells [1]. Type I cells are thought to primarily function as support cells and share some characteristics of glia cells [2], while Type II cells detect bitter, sweet or umami stimuli through the activation of specific GPCRs for each of these taste qualities. Transduction of these taste qualities in Type II cells is due to a single signaling pathway that is comprised of phospholipase Cβ2 (PLCβ2) which activates the inositol 1,4,5-trisphosphate receptor type 3 (IP_3_R3) on the endoplasmic reticulum to cause calcium (Ca^2+^) release [3–5]. This Ca^2+^ release activates the transient receptor potential cation channel subfamily M members 4 and 5 (TRPM4 and TRPM5) which depolarize the cell sufficiently to activate the release of ATP through the calcium homeostasis modulator 1 (CALHM1) channel [4, 6–10]. The expression of PLCβ2, IP_3_R3 and TRPM5 is restricted to Type II taste cells and these cells lack both voltage-gated Ca^2+^ channels (VGCCs) and conventional chemical synapses (model shown in Figure 1A)[11–15]. Type III cells detect sour and salt stimuli through ionotropic receptors that depolarize the cell to activate VGCCs and cause vesicular neurotransmitter release (see Figure 1A) [16–22]. It is currently thought that Type III cells do not respond to bitter, sweet or umami stimuli.

**Figure 1.**
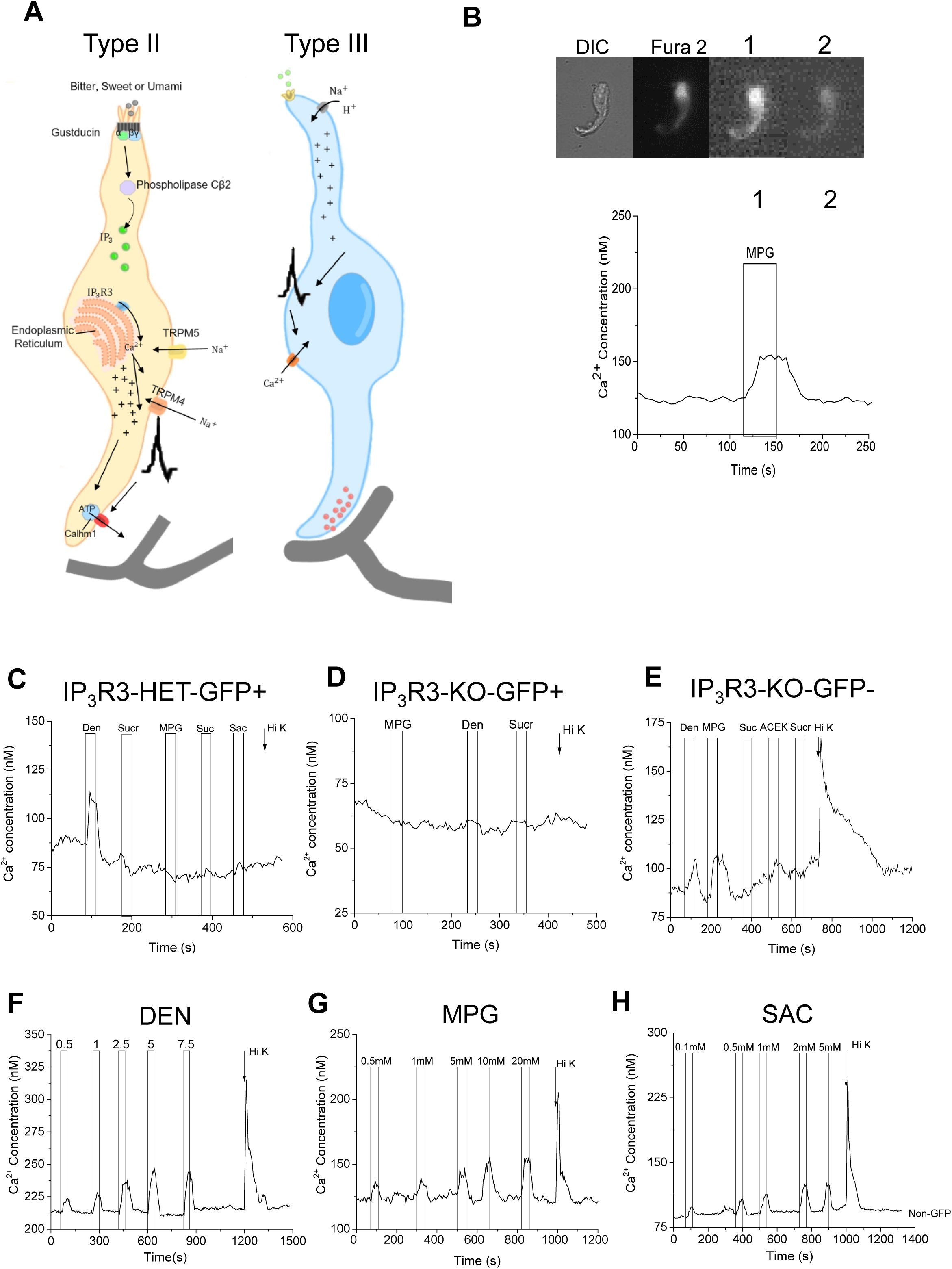
Current taste cell models and experimental setup. A) Current models of the signal transduction pathways in Type II taste cells (left) that respond to bitter, sweet, and umami stimuli as well as Type III taste cells (right) that respond to sour and salty stimuli. B) Example of an isolated taste cell that was stimulated with MPG (10mM). The DIC image is shown first, followed by an image of Fura 2-AM loading. The next two images show the ratio (340/380) at different time points as the cell is stimulated and then recovers to baseline. The corresponding calcium (Ca^2+^) changes are shown in the imaging trace. C) The IP_3_R3-HET-GFP+ cell responded to denatonium, but did not respond to any of the other stimuli applied. D) The IP_3_R3-KO-GFP+ cell did not respond to any stimulus applied. E) The IP_3_R3-KO-GFP-cell responded to denatonium, MPG and 50mM KCl (denoted with arrow, Hi K), identifying it as a BR cell. Concentration gradients for (F) bitter (denatonium, DEN), (G) umami (MPG) and (H) sweet (saccharin, SAC). BR cells respond to a wide range of stimulus concentrations.

We previously reported that some taste cells generate Ca^2+^ signals to both denatonium, a bitter stimulus and cell depolarization with 50mM KCl [23]. Since cells expressing VGCCs respond to cell depolarization with a Ca^2+^ signal, these data identified a potentially unique taste cell population that expresses VGCCs but also responds to denatonium [23]. To date, these cells have not been characterized. We hypothesized that there is a subset of Type III taste cells that are responsive to denatonium and are potentially sensitive to multiple bitter, sweet and umami stimuli. To address this question, we used live cell imaging in an IP_3_R3-KO mouse line to evaluate these cells without Ca^2+^ responses from Type II cells. We identified a subset of Type III cells that respond to bitter, sweet and/or umami stimuli and determined that these broadly-responsive (BR) taste cells use a PLCβ3 signaling pathway. The loss of PLCβ3 in the BR cells caused significant deficits in subsequent taste perception, supporting the conclusion that these BR cells are important for taste.

## RESULTS

### Bitter, sweet and umami taste evoked responses are still present in IP_3_R3-KO mice

To assess the characteristics of the BR cells, we wished to avoid any potential signaling input from the Type II cells that respond to bitter, sweet and umami stimuli. IP_3_R3 is part of the canonical signaling pathway in Type II cells that are sensitive to these stimuli [3, 11, 15, 24]. Therefore, we used a transgenic mouse in which GFP replaces the coding region of IP_3_R3 to evaluate the taste evoked signaling in taste receptor cells that lack the ability to release Ca^2+^ via IP_3_R3. In these mice, GFP labels the cells that should express IP_3_R3 but no longer do so. Initial immunohistochemical analyses found that taste cells from the CV papillae of wild type mice were successfully labeled with anti-IP_3_R3 antibody (n=3; Figure S1A), while their KO littermates lacked IP_3_R3 labeling but instead had GFP expression (n=6; Figure S1B). Furthermore, anti-IP_3_R3 labels the GFP expressing cells in IP_3_R3-heterzygous mice (n=4; Figure S1C); confirming that GFP expression identifies the taste cells that normally express IP_3_R3. To further characterize this mouse, we evaluated the expression of other proteins that are part of this established signaling pathway in Type II cells. The GFP expression in the KO mice co-localized with both PLCβ2 (n=4; Figure S1D) and gustducin (n=3; Figure S1E) labeling. As expected, gustducin expression was restricted to the IP_3_R3-GFP-KO cells but not all GFP+ cells had gustducin. There was, however, complete overlap between anti-PLCβ2 labeling and GFP. Co-localization analyses are reported in Figures S1F. Thus the other components of the signaling pathway normally expressed in Type II cells are intact but these cells no longer have the capacity to release Ca^2+^ in response to bitter, sweet or umami stimuli.

We also functionally characterized the ability of the taste cells in this mouse to respond to taste stimuli. Measurements were made in isolated individual taste cells to ensure that the measured taste-evoked Ca^2+^ responses were due solely to the activity of the individual taste cells and not cell-to-cell communication (example of a response in an isolated cell in Figure 1B). To functionally characterize the IP_3_R3-GFP-KO mouse, and to confirm that GFP is expressed in Type II cells, we measured the taste-evoked Ca^2+^ responses in GFP+ taste cells from the IP_3_R3-het mouse which express GFP but also have a functional copy of the IP3R3 gene. These cells responded to taste stimuli but did not generate a Ca^2+^ signal to 50mM KCl, indicating these cells do not express VGCCs (Figure 1C) and are likely Type II cells. Alternatively, GFP+ cells from the IP_3_R3-KO mouse (lack a functional IP_3_R3 gene) did not produce Ca^2+^ signals when stimulated (Figure 1D). When we targeted the GFP-cells in the IP_3_R3-KO mouse, we identified taste cells that responded to both taste stimuli and 50mM KCl with a Ca^2+^ signal (Figure 1E). These data suggest that these BR taste cells do not require input from Type II cells to respond to bitter, sweet or umami stimuli.

We characterized BR cells by applying taste stimuli to isolated taste cells from IP_3_R3-KO mice. Since our goal was to identify all (or most) responsive taste cells, we used the stimulus concentration that generated a maximal Ca^2+^ signal. For all experiments, taste stimuli were applied for 40s (application shown with bar in graphs) and 50mM KCl was applied for 10s (application shown with arrow in graphs). These concentrations were based on our previous control experiments to identify the lowest stimulus concentration that generated the maximal Ca^2+^ signal (examples shown in Figure 1F-H).

Type III cells are the only known population of taste cells that express VGCCs [25] and are functionally identified in live cell imaging by the ability to respond to cell depolarization with a Ca^2+^ influx through the opening of VGCCs [15, 18, 25–28]. Therefore, if a taste cell responded to 50mM KCl with a Ca^2+^ signal, we concluded that it expresses VGCCs and is a Type III cell. We applied multiple bitter, sweet and umami stimuli + 50mM KCl to isolated taste receptor cells from IP_3_R3-KO mice (Figure 2A-C). We found that BR cells often responded to multiple stimuli, including multiple types of stimuli for each taste quality. To evaluate if the loss of IP_3_R3 affected this population of BR cells, we compared the percentage of BR cells between the WT (n=105 mice) and IP_3_R3-KO mice (n=115). Because taste-evoked response sizes are variable, we focused our analysis on the frequency of cell responses. A response was recorded if the isolated taste cell had a stable baseline and application of the taste stimulus caused an increase in fluorescence that was at least 2 standard deviations above the baseline fluorescence level. We found no significant differences in the frequency of taste cells that were BR in the IP_3_R3-KO mice compared to WT (Figure 2D). This data indicates that the BR cell responses do not rely on Type II cell responses.

**Figure 2.**
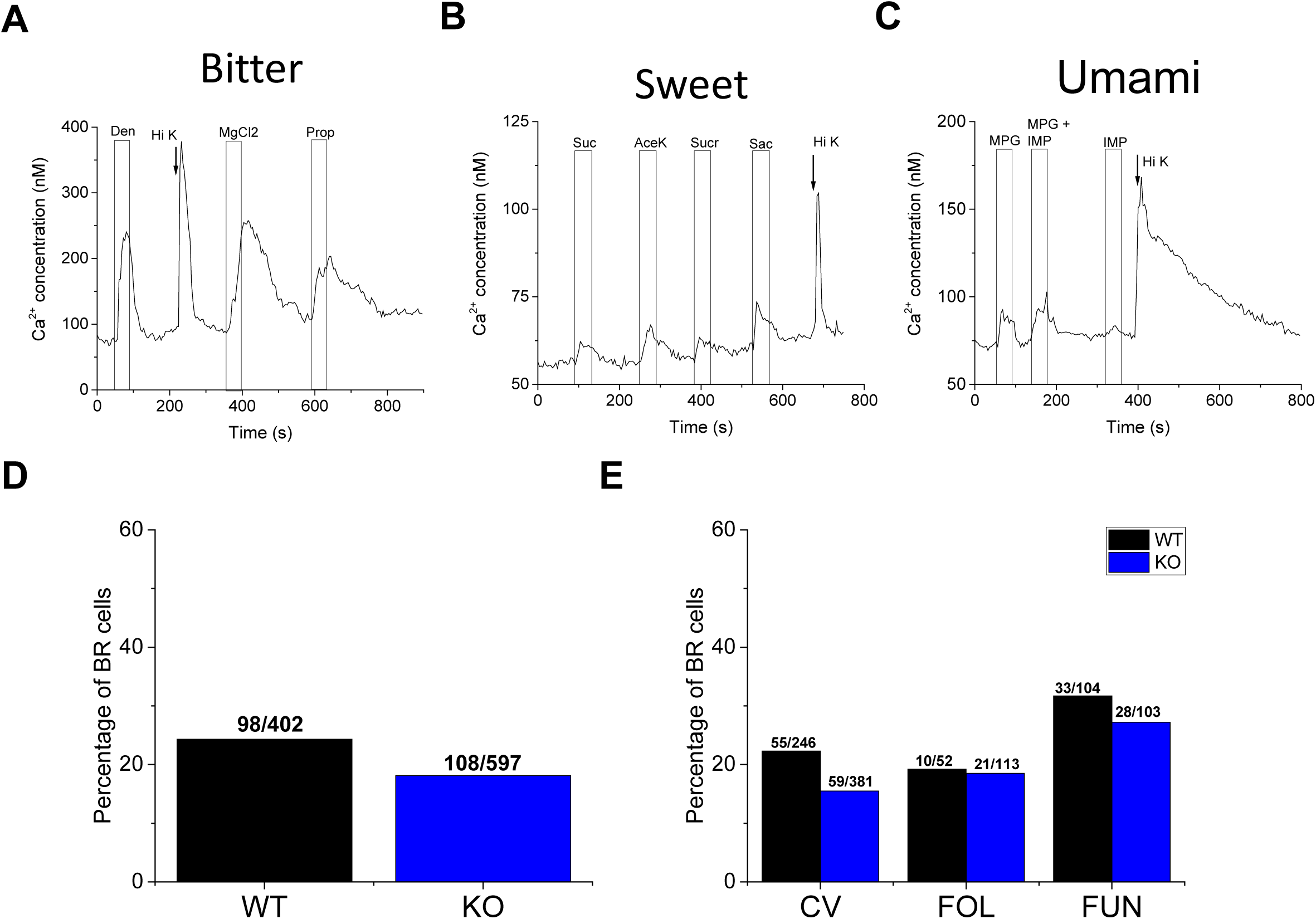
BR taste cells respond to multiple taste stimuli independently of the IP_3_R3 protein expressed in Type II cells. Representative traces of BR taste cells that were stimulated with multiple bitter (A), sweet (B), or umami (C) stimuli as well as 50mM KCl (arrow, Hi K). D) Chi square analysis with Yate’s correction for continuity was used to compare the frequency of evoked Ca^2+^ responses to different taste stimuli between wild type (WT) and IP_3_R3-KO (KO) mice (p=0.062). E) Chi square analysis with Yate’s correction for continuity was used to compare the frequency of evoked Ca^2+^ responses by BR cells from wild type (WT) and IP_3_R3-KO (KO) mice for circumvallate (CV, p=0.09), foliate (Fol,p=0.89) and fungiform (Fun,p=0.7) taste cells.

There are three papillae types that house taste buds in different areas of the tongue: circumvallate (CV), foliate (Fol), and fungiform (Fun). We measured the response frequency of BR cells for the different papillae types in the WT and IP_3_R3-KO mice and found no significant differences (Figure 2E). Thus, BR cells are present in multiple taste papillae and the loss of IP_3_R3 in Type II cells does not affect the percentage of taste cells that are BR.

### A specific subset of Type III cells is broadly responsive (BR)

The current understanding is that all Type III cells are sour sensitive and a subset of these sour-sensitive Type III cells are also salt sensitive [16, 18, 28]. To confirm that the BR taste cells are Type III cells, we included sour (citric acid, CA) and salt (NaCl) stimuli in some experiments (Figure 3). In these experiments, we used a representative stimulus for bitter (5mM Den), sweet (20mM sucralose) and umami (10mM MPG). All of the taste cells that responded to 50mM KCl responded to citric acid (CA) with a Ca^2+^ signal while some of these cells also responded to bitter (B), sweet (S) and/or umami (U) stimuli with a Ca^2+^ signal (Figure 3A, C, D). A representative imaging trace is shown in Figure 3A, with the summary of the Type III responses shown in Figure 3C. We did not detect any Type III cells that responded to all 5 taste qualities (Figure 3B, C, orange region). However, in addition to the Type III cells that responded to either sour (CA, red region) or sour and salty (CA + NaCl, blue region) there was another subset of Type III cells (52%, yellow region) that responded to sour and at least one bitter, sweet, and/or umami (B, U, S) stimulus. To better understand these BR cells, we increased our sample size and measured the distribution of the taste responses to bitter, sweet and umami stimuli within these cells (Figure 3D). Unlike the Type II cells, few BR cells responded to a single taste quality (bitter, sweet and umami were 5.5, 7 and 5.5% respectively, blue regions) but were instead most likely to respond to two or three taste qualities in addition to responding to 50 mM KCl. 39% of the cells responded to two stimuli (green region) while 43% responded to all three (gray region).

**Figure 3.**
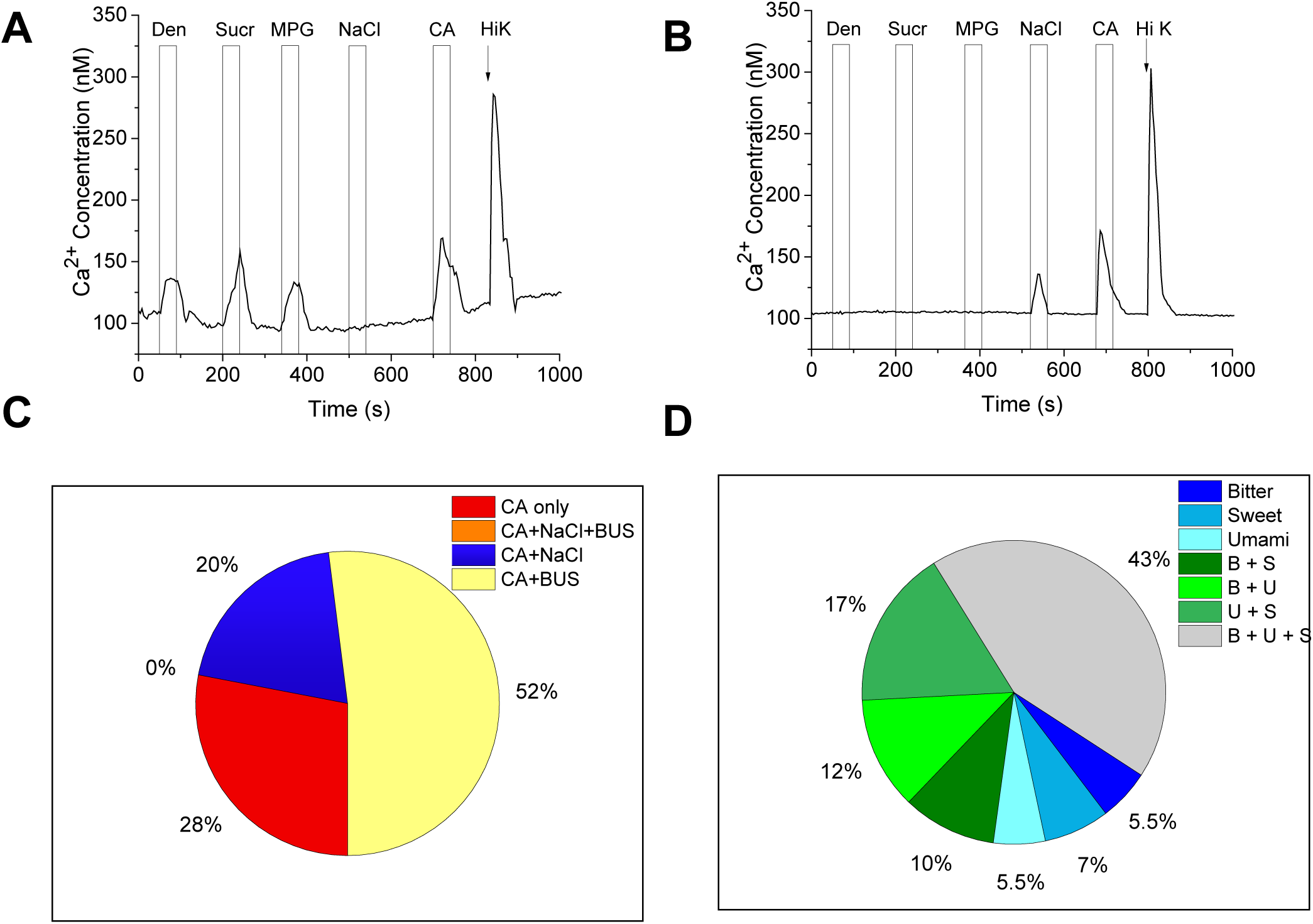
Some Type III cells respond to bitter, sweet and umami stimuli. A) Representative trace of taste cells from IP_3_R3-KO mice that responded to bitter (5mM denatonium, Den), sweet (20mM sucralose, Sucr), umami (10mM monopotassium glutamate, MPG), 50mM citric acid (CA), and 50mM KCl (Hi K). B) Representative trace of a separate subset of Type III cells that responded to 250mM NaCl, 50mM CA and 50mM KCl but were not sensitive to the bitter, sweet, umami stimuli tested. C) Summary of taste cells from IP_3_R3-KO mice that responded to 50mM KCl and CA with Ca^2+^ signals (n=65), 28% only responded to CA (n=18), 20% responded to CA and NaCl (n=13), and 52% responded to CA and bitter, sweet and/or umami stimuli (B,U,S, n=34) while no cells responded to all taste stimuli. D) A summary of the response profiles for a larger group of BR Type III cells (n=127) that either responded to bitter (n=7), sweet (n=9), umami (n=7), bitter + sweet (n=13), bitter + umami (n=15), sweet + umami (n=21) or bitter, sweet + umami (n=55).

Control experiments using the GAD67-GFP mice as a cellular marker to identify Type III taste cells [29–31] found that both GAD67-GFP positive and negative taste cells responded to taste stimuli + 50mM KCl with Ca^2+^ signals (Figure S2A-B). Approximately 69% (n=98/143 cells tested) of the GAD67-GFP positive cells responded to a bitter, sweet or umami stimulus as well as 50mM KCl. BR cells were also identified in C57BL/6 mice (Figure S2C) which agrees with a separate study that reported some Type III cells from C57BL/6 mice responded to bitter stimuli [18]. Together these data suggest that these BR Type III taste cells are present in taste cells in multiple mouse lines.

### BR cells use a PLCβ signaling pathway to respond to bitter, sweet, and/or umami stimuli

To identify the type of signaling pathway that generates the taste responses in the BR cells, we analyzed the amplitudes of the responses to bitter (5mM Den), sweet (20mM sucralose) and umami (10mM MPG) stimuli in the IP_3_R3-KO mouse. We found that the size of the responses to these stimuli did not significantly change in Ca^2+^-free Tyrode’s, indicating these signals depend on Ca^2+^ release from internal stores (Figure 4A, D, G). We confirmed these results by applying thapsigargin to disrupt the internal Ca^2+^ stores which abolished the taste evoked Ca^2+^ signals (Figure 4B, E, H). These responses were also eliminated when the general PLC blocker U73122 was applied (Figure 4C, F, I), indicating that these taste evoked Ca^2+^ responses in the BR are due to the activation of a PLC signaling pathway. Representative traces for each experimental condition are shown in Figure S3.

**Figure 4.**
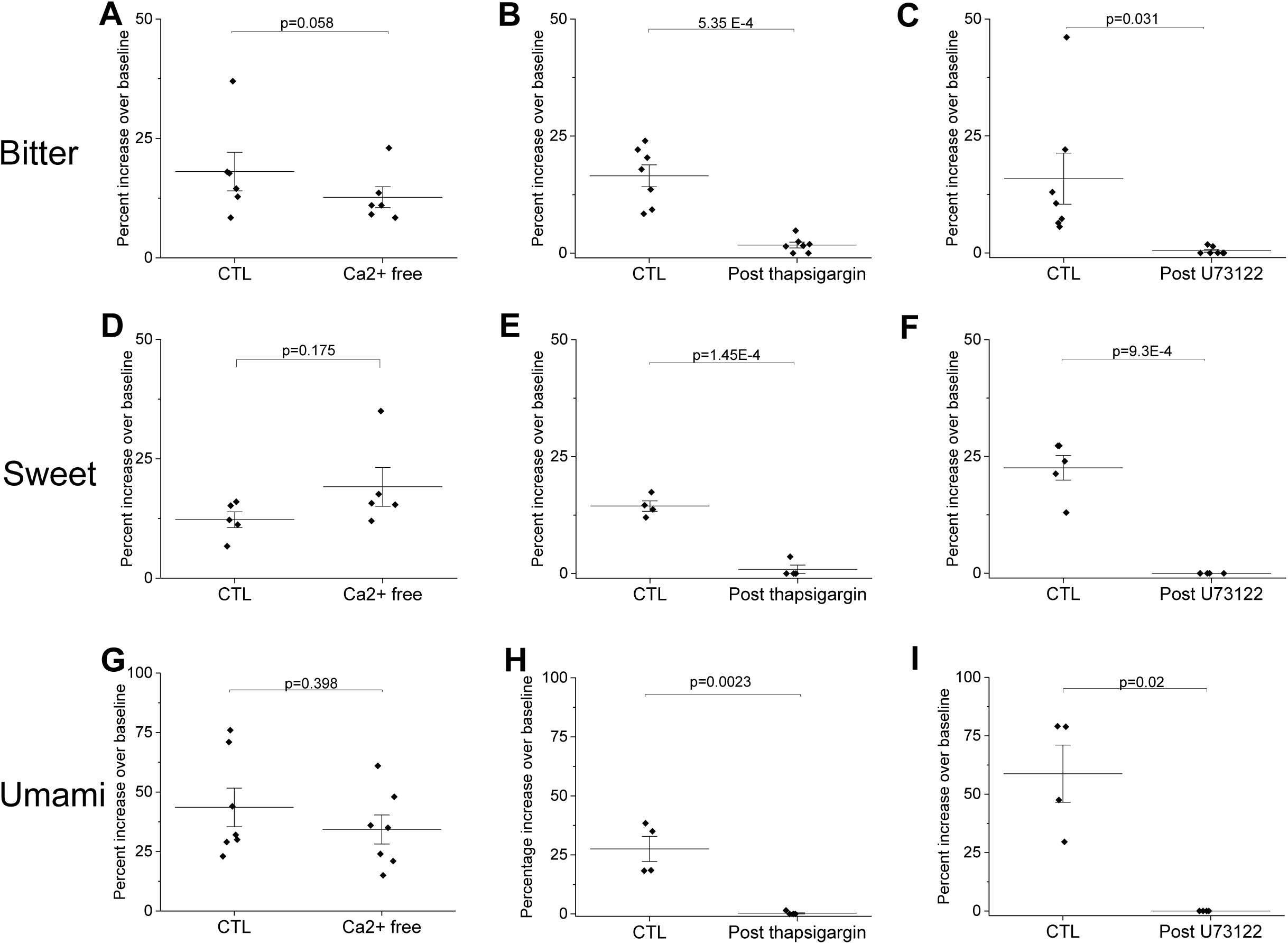
Taste-evoked Ca^2+^ release in BR cells from IP_3_R3-KO mice is dependent upon PLC activity and Ca^2+^ release from internal stores. A) Bitter-evoked taste responses (5mM Denatonium, n=6) persist in the absence of extracellular calcium (Ca^2+^-free) and are abolished by the SERCA pump inhibitor thapsigargin (B, n=7) as well as the PLC blocker U73122 (C, n=7). D) Responses to sweet stimuli (50mM sucrose, n=6) persist in Ca^2+^-free and are abolished by thapsigargin (E, n=4) and U73122 (F, n=6). G) Umami stimuli (10mM MPG) persist in Ca^2+^-free (n=7) and were abolished by thapsigargin (H, n=4) and U73122 (I, n=4). Representative data for each experiment are shown in Figure S3. Comparisons of the response amplitudes for each taste stimulus found no significant differences in the control responses between experiments (One way ANOVA, Den, p=0.933; Suc, p=0.623; MPG, p=0.134).

### PLCβ3 expression in peripheral taste cells

To identify other PLCβ isoforms expressed in taste receptor cells, we analyzed our previously published RNAseq data [32]. We found that PLCβ2 and PLCβ3 were expressed at comparable levels and were the predominant PLCβ isoforms present in taste cells. Since PLCβ2 is only expressed in Type II cells, we investigated the possibility that PLCβ3 is expressed in peripheral taste cells. Immunohistochemical experiments in the IP_3_R3-KO mouse revealed strong PLCβ3 labeling that was separate from the Type II cells (identified with GFP expression, Figure S4A). These data were confirmed by examining anti-PLCβ3 labeling in the TRPM5-GFP mouse as well as co-labeling with PLCβ2 in C57BL/6 mice (additional Type II cell markers, Figure S4B, C). Co-labeling with the Type I taste marker NTPDase2 found that PLCβ3 is not expressed in Type I cells (Figure S4D). We next determined if PLCβ3 is expressed in Type III cells using anti-SNAP-25 which is a synaptic protein that is a cellular marker for Type III taste cells [15, 29, 33]. Immunohistochemical analysis found some co-expression of PLCβ3 and SNAP25 in C57BL/6 mice (Figure S4E) supporting the conclusion that PLCβ3 is expressed in some Type III cells. Colocalization analyses are shown in S4F.

We also used qPCR to measure the relative amounts of mRNA for PLCβ3 and PLCβ2 in the different taste papillae (Figure S4G, H) and found that both PLCβ2 and PLCβ3 are expressed in CV, Fol and Fun papillae. Therefore, both PLCβ2 and PLCβ3 are expressed in peripheral taste cells and PLCβ3 is expressed in a subset of Type III cells but not either Type I or Type II taste cells.

### PLCβ3 contributes to taste evoked signaling in a subset of Type III cells

We next performed live cell imaging on isolated taste receptor cells from PLCβ3-KO and wild type mice to evaluate how the loss of PLCβ3 affects the responsiveness of the BR cells. Type III cells in the PLCβ3-KO mouse were still present and responded to sour and salt stimuli (Figure 5A, B). A summary of the response profiles is shown in Figure 5C. Some Type III cells responded to sour (CA, red region) or sour + NaCl (blue region), but none responded to any bitter, sweet, or umami stimuli (yellow region) tested. Type II cells were still functional and responded to these stimuli (black region). The responsiveness of Type II cells was not significantly affected by the loss of PLCβ3 expression (Figure S5). Example traces of Type II responses are shown in Figure 5D-F. A more comprehensive analysis of the PLCβ3-KO mice found that the loss of PLCβ3 caused a very significant reduction in the number of BR taste cells (Figure 5G) and no BR cells were identified in either the Fol or Fun papillae of the PLCβ3-KO mouse (Figure 5H). While 23.5% of the CV taste cells were BR in the WT mice, only 2% of the CV taste cells were BR in the PLCβ3-KO mouse. Immunohistochemical analysis of the PLCβ3-KO mouse confirmed the absence of any PLCβ3 expression in the taste buds (Figure S6) compared to WT mice. These data suggest that PLCβ3 is required for transducing most of the taste evoked responses in BR cells.

**Figure 5.**
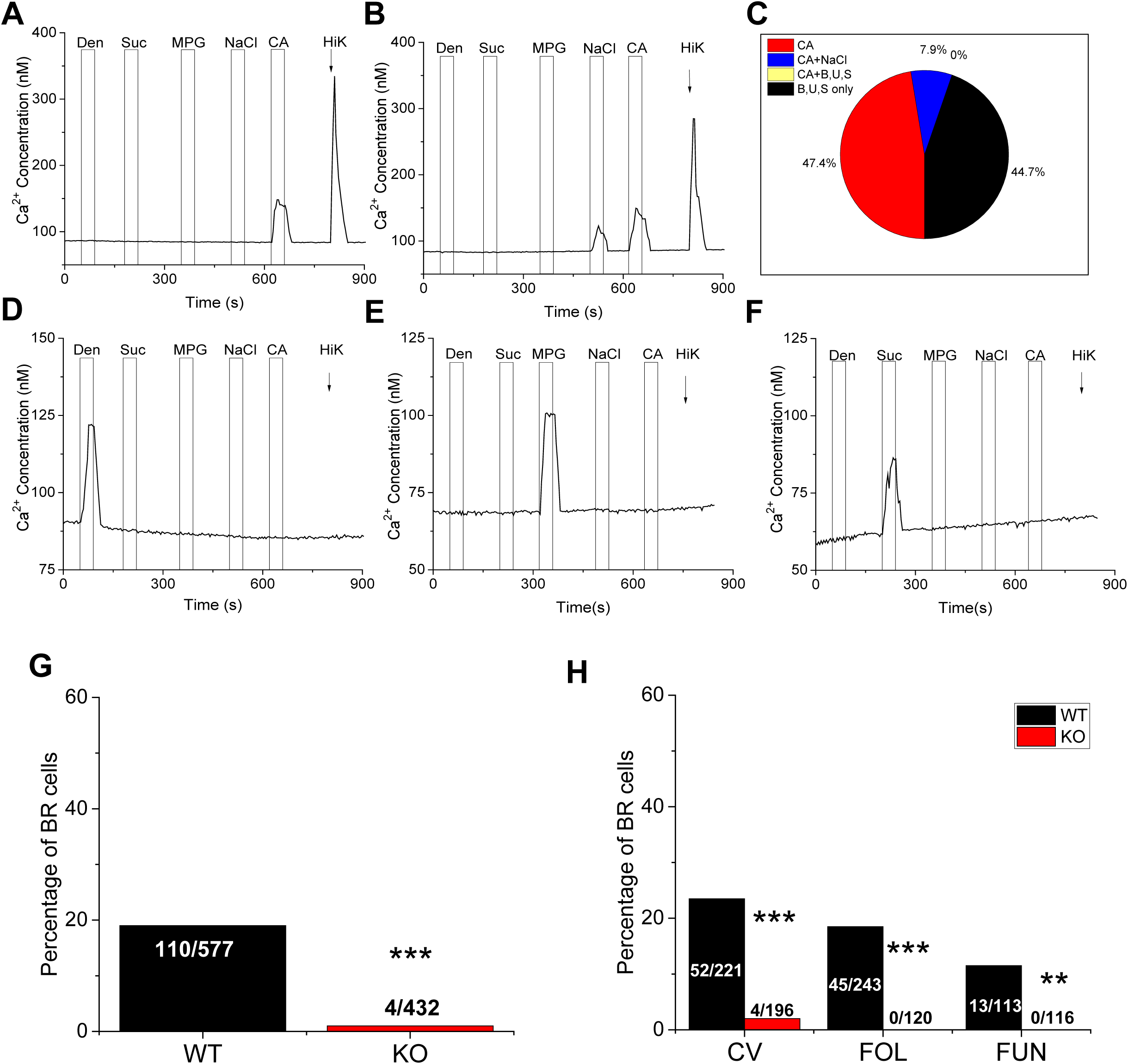
PLCβ3 is required for the detection of bitter, sweet and umami stimuli in BR cells. A) A representative trace from PLCβ3-KO taste cells that responded to 50mM citric acid (CA) and KCl (50mM, Hi K) but did not respond to bitter (5mM denatonium, Den), sweet (20mM sucralose, Sucr) or umami (10mM MPG) stimuli. B) A representative trace from a separate subset of cells that responded to 250mM NaCl, CA and KCl but were not sensitive to the bitter, sweet, umami stimuli tested. C) Summary of responsive taste cells from PLCβ3-KO mice (n=38): CA (n=18); CA +NaCl (n=3); CA and bitter, sweet and/or umami stimuli (B, U, S) (n=0); and B,U, or S only (n=17). Representative taste-evoked responses for bitter (D), sweet (E) and umami (F) in the PLCβ3-KO mice. None of these cells responded to CA or KCl. G) Chi square analysis with Yate’s correction for continuity was used to compare the frequency of BR cells in wild type (WT, n=53 mice) and PLCβ3-KO (KO, n=68) mice (***, p=0) for a larger number of cells. H) The percentage of BR taste cells from the CV, Fol and Fun papillae of the PLCβ3-KO mice and WT mice were compared (***, p<0.001 for CV and Fol; **, p=0011 for Fun).

### Loss of PLCβ3 causes deficits in the taste-evoked activity in the nucleus of the solitary tract

Our data so far demonstrate that a subset of Type III taste cells are capable of responding to bitter, sweet and umami using a PLCβ3 signaling pathway. To determine if these BR taste cells are important in taste transduction, we measured the effect of BR cells on the taste signal that is sent to the brain. This was accomplished by measuring the neural activity in the nucleus of the solitary tract (NTS) using c-Fos labeling as a marker for activated neurons. The NTS receives input from the gustatory neurons and is the first synaptic relay from the peripheral taste system. This approach is commonly used to identify recently activated neurons, including neurons in the taste pathway [34–38]. We reasoned that if BR cells are contributing to the transduction of taste information to the brain, then the loss of PLCβ3 would have a significant impact on the level of c-Fos labeling in the NTS. We used the IP_3_R3-KO mouse as a positive control since it has previously been shown to be required to transmit bitter, sweet and umami taste information to the brain [39]. Mice (n=3 for each mouse line) were orally infused with quinine (5mM) or water for 30 min. Background c-Fos labeling was measured using water as a control. As expected, WT mice had a large number of c-Fos positive cells in the NTS (Figure 6A, upper left panel), while the IP_3_R3-KO mice (Figure 6A, upper right panel) and water control mice (Figure 6A, lower right panel) both had few c-Fos positive cells. Strikingly, the level of c-Fos labeling in the PLCβ3-KO mice (Figure 6A, lower left panel) were comparable the c-Fos levels in the IP_3_R3-KO mice. Thus, loss of either of these proteins resulted in a significant reduction in c-Fos activity to a level that was comparable to water (Figure 6B). Control experiments demonstrate that neither IP_3_R3 nor PLCβ3 are expressed in this region of the NTS (Figure S7) so the recorded deficits were not due to the loss of either of these proteins in the NTS.

**Figure 6.**
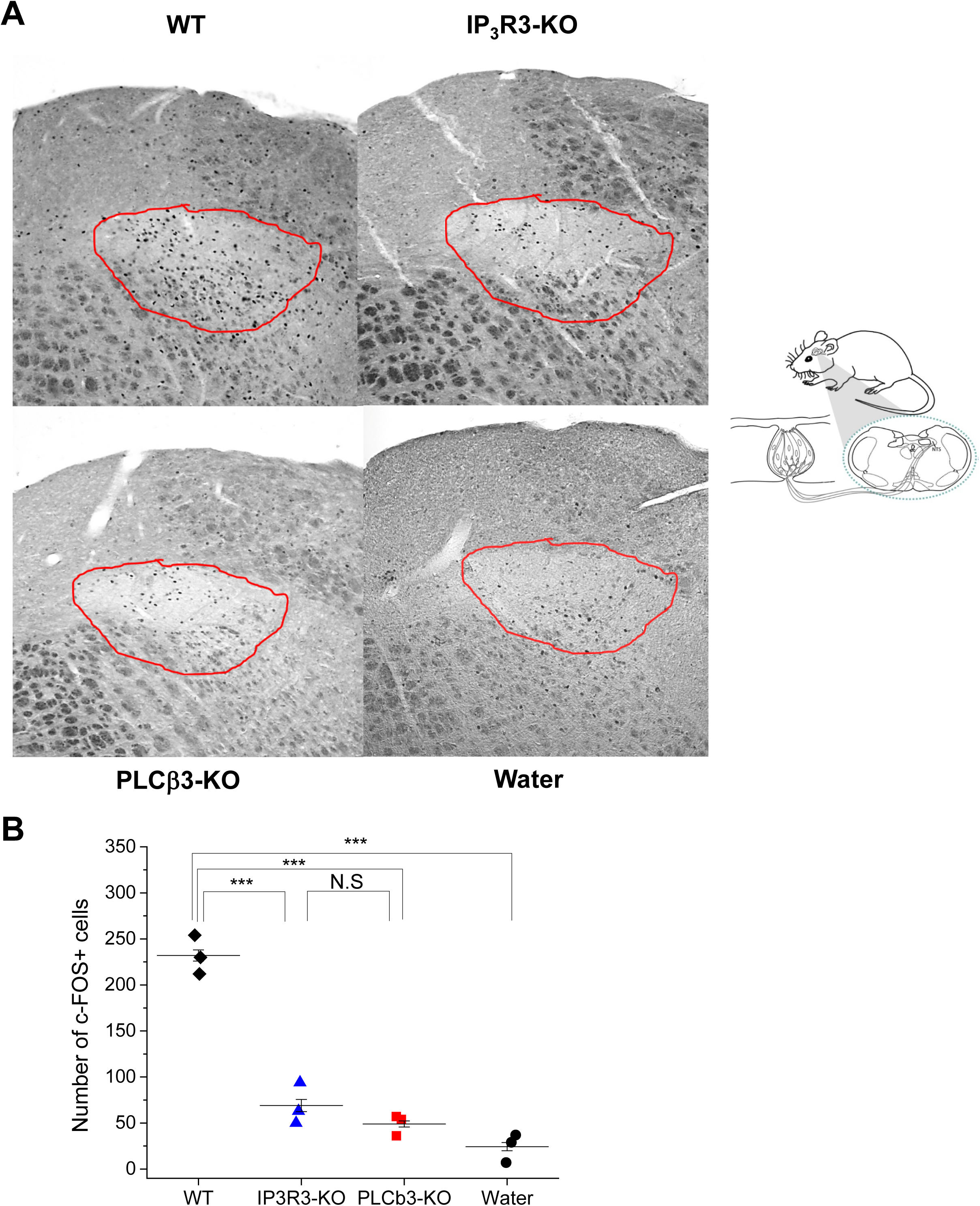
Loss of PLCβ3 or IP_3_R3 causes deficits in the taste-evoked neural activity in the nucleus of solitary tract. Oral infusion of bitter (quinine) and water elicited c-Fos immunoreactivity in the intermediate rostral nucleus tractus solitarius (IRNTS). The schematic diagram indicates the region in the hindbrain that was analyzed. (A, B) Loss of either PLCβ3 (PLCβ3-KO) or IP_3_R3 (IP_3_R3-KO) caused a significant reduction in the c-Fos immunoreactivity in the IRNTS compared to WT mice. The residual c-Fos labeling in each of the KO mice was comparable to the c-Fos labelling generated by water infusion. For all experiments, 3 mice of each genotype were used (***, p<0.001; **, p<0.01). The red outlined area on the DIC images denotes the analyzed region for each. Data were analyzed with a one way ANOVA with follow up Student’s t tests to identify individual differences. Significance level was set at p<0.05.

### Loss of PLCβ3 significantly affects taste-driven licking for bitter, sweet and umami

Since our data indicate that the BR cells are contributing to the taste signals that are sent to the brain, we then performed brief-access licking experiments to determine if the loss of taste-evoked signals in the PLCβ3-KO mice correlated with loss of taste sensitivity at the behavioral level. Measurements were also made with IP_3_R3-KO mice for comparison. Since the PLCβ3-KO mouse has a mixed genetic background while the IP_3_R3-KO mice are in a C57BL/6 background, wild type littermates for both mouse lines were used as controls (Figure 7). The wild type mice both demonstrated concentration dependent decreases in licking to denatonium while PLCβ3-KO and IP_3_R3-KO mice treated the solution similarly to water except for the highest concentration (20mM Den, Figure 7A). Wild type mice treated MSG (+ 10µM amiloride) solutions as more palatable than water up to 400mM and then decreased licking at the highest concentrations (600 and 800mM, Figure 7B). Both of the KO mice treated MSG like water. In the brief access test for the artificial sweetener, Acesulfame K (AceK), wild type mice increased licking up to 2 mM and then decreased licking at higher concentrations. Both of the KO mice treated AceK like water (Figure 7C). Because sucrose is hedonically positive under wild type conditions, the lick score is calculated as licks to the stimulus minus licks to water, therefore a score close to zero resembles water. Neither the PLCβ3-KO nor IP_3_R3-KO mice showed a concentration-dependent effect on licking to sucrose until they were presented with the highest concentration. In contrast, both sets of wild type mice showed strong concentration dependent increases in licking to sucrose (Figure 7D). As a positive control, we also tested sodium chloride since salt is not predicted to activate a PLCβ signaling pathway in taste cells. For the salt stimulus (NaCl), there were no significant differences in the licking behavior for either of the KO mice compared to wild type mice, indicating that the ability to respond to salt is not affected by loss of either PLCβ3 or IP_3_R3 (Figure 7E). Thus, loss of either IP_3_R3 or PLCβ3 caused comparable, and almost complete, loss of behavioral responses specifically for bitter, sweet and umami stimuli.

**Figure 7.**
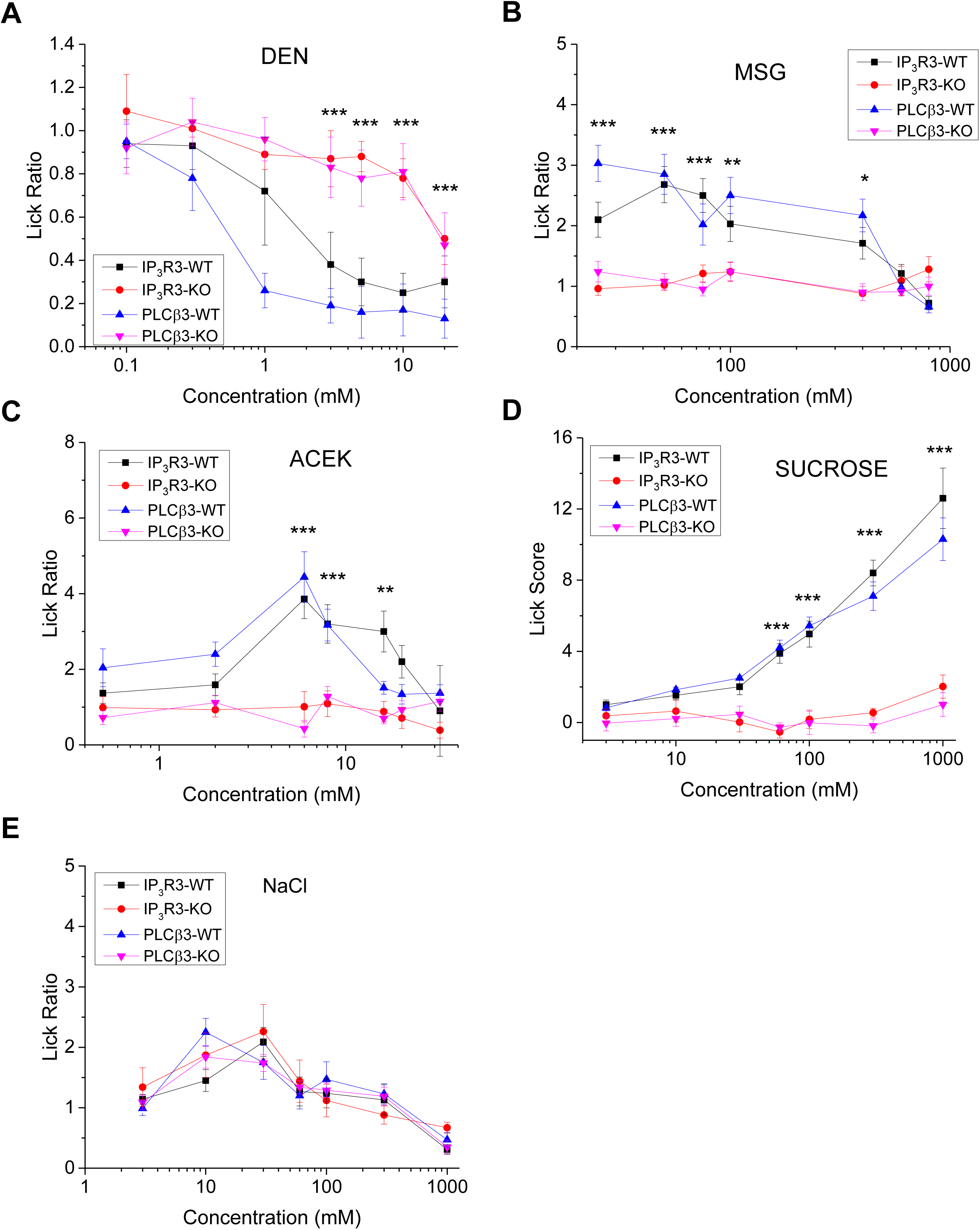
Loss of PLCβ3 or IP_3_R3 affects behavioral responses to taste stimuli. Average lick data (±standard deviation) from brief-access behavioral tests compare the responses of IP_3_R3-KO (red line) and PLCβ3-KO (pink line) to WT (IP_3_R3-WT, black line; PLCβ3-WT, blue line). A) Lick ratios (stimulus/water) of the WT mice for the bitter stimulus denatonium (DEN, 0, 0.1, 0.3, 1, 3, 5, 10, 20mM) were significantly different from the IP_3_R3-KO and PLCβ3-KO responses. B) Lick ratios of the WT mice for the umami stimulus monosodium glutamate + 10µM amiloride (MSG, 0, 25, 50, 75, 100, 400, 600, 800mM) were significantly different from the IP_3_R3-KO and PLCβ3-KO mice. C) Lick ratios of the WT mice for the artificial sweetener acesulfame K (ACEK, 0, 0.5, 2, 6, 8, 16, 20, 32mM) were significantly different from the IP_3_R3-KO and PLCβ3-KO mice. D) Lick scores (stimulus-water) of the WT mice for sucrose (0, 3, 10, 30, 60,100, 300, 1000mM) were significantly different from the IP_3_R3-KO and PLCβ3-KO responses. E) No significant differences were detected between the WT and KO mice for any concentration of NaCl (0, 3, 10, 30, 60, 100, 300, 1000mM) tested. For all experiments, 5 mice of each genotype were used (***, p<0.001; **, p<0.01; *, p<0.05). Data were compared by repeated measures ANOVA. Significant interaction terms were followed by Tukey’s Honestly Significant Difference tests.

## DISCUSSION

This study has characterized a population of taste cells that express VGCCs, respond to sour stimulation with a Ca^2+^ signal, but are also responsive to bitter, sweet and/or umami stimuli. Our data suggest that BR cells are a subset of Type III cells that are capable of responding to multiple taste stimuli, except sodium chloride. Since the BR cells always responded to sour, 100% of these cells responded to multiple taste qualities and approximately 80% of these cells responded to either three or four modalities (Figure 3D). We also found that the BR cells could respond to multiple stimuli within the same quality. This is in contrast to Type II cells that are usually narrowly tuned to a single taste quality. While we are not able to stimulate individual BR cells with all taste stimuli, our data suggest that the BR cells may be very broadly tuned and act as a generalist responding cell. It is unclear why BR cells did not respond to the sodium salt we tested, but other studies have reported that sodium responsive cells appear to be a separate taste cell population [18, 40].

The idea of broadly tuned taste cells in mammals has been put forth by multiple labs, using a variety of approaches that utilized whole taste buds [41–48]. Maintaining the taste cells in the bud allowed for measurements in a more “native” environment and allowed for stimuli to be apically applied to the cells. Sato and Beidler [42, 43] inserted a microelectrode into a taste cell through the taste pore, thus leaving the taste bud intact within the tongue of an anesthetized rat. Using this minimally invasive approach, they identified that most taste cells (∼85%) responded to more than one type of taste stimuli. Similarly, Gilbertson et al [48] apically applied four basic taste stimuli to taste buds within the epithelium using a modified Ussing chamber. Their approach allowed for the focal application of taste stimuli to the apical end of the taste cells that were still within the bud. Using patch clamp analysis, they found approximately 73% of the cells responded to two or more taste stimuli [48]. Ninomiya’s group analyzed Type II and Type III taste cells in excised mouse taste buds using gustducin-GFP to identify Type II cells and GAD67-GFP to identify Type III cells [31]. They found that Type II taste cells (gustducin-GFP+) were selective in their responses to bitter, sweet or umami stimuli while all Type III cells (GAD67-GFP+) responded to sour stimuli. Interestingly, they found about 25% of the GAD67-GFP+ cells responded to multiple taste stimuli, including sweet, bitter and umami in addition to sour [31]. Roper’s lab used calcium imaging in a slice preparation of taste buds in which they focally applied taste stimuli [30, 47] and also found that a large percentage of the taste cells were broadly tuned. These studies also concluded that Type II cells are more selective in their responsiveness while Type III cells are sometimes broadly tuned [30].

All of these earlier studies used intact taste buds and in some cases, the authors concluded that the broadly responsive Type III cells were likely receiving input from the neighboring Type II cells [30]. A key difference between our work and these earlier studies is that we used isolated taste cells that were not in contact with neighboring cells. This allowed us to analyze how individual taste cells respond without any potential input from neighboring cells. While we do not rule out the possibility that taste cells receive input from other taste cells, our data show that BR Type III cells do not require neighboring input to respond to bitter, sweet and/or umami stimuli. Taken with these earlier studies [30, 31, 41–48], our data support the idea that there are both selectively and broadly responsive cells in taste buds. In agreement with others [30, 31], our data indicate that these BR cells are a subset of Type III taste cells and we have now shown that BR cells can independently respond to bitter, sweet and umami stimuli.

Many studies focused on the upstream components of the taste pathway have also reported that generalist and specialist cell populations exist. Studies of individual afferent gustatory neurons and geniculate ganglion cells found that some gustatory nerves are highly specific to a particular taste stimulus (specialists) while many of these cells are broadly tuned (generalists) [48–54]. At the first synaptic relay of central taste processing, some neurons in the rostral nucleus of the solitary tract (rNTS) respond ‘best’ to a particular taste stimulus, while others are broadly tuned [55, 56]. Specialists and generalists cells have also been described in the gustatory cortex [57–61]. Our data demonstrate that taste receptor cells, the first step in the taste pathway, mirror the functional heterogeneity that is found higher up in the taste processing; some taste cells are selective (Type II taste cells) to a particular stimuli, while others are broadly tuned (BR taste cells) to multiple taste stimuli.

While we previously reported that PLCβ3 is expressed in a subset of taste cells that are not Type II cells [23], we have now demonstrated PLCβ3 is present in a subset of Type III cells. This idea is supported by recent RNA-sequencing analysis of individual Type II and Type III cells [26] which identifies PLCβ3 expression in 30% of Type III cells. We searched this RNA-seq dataset of Type III cells [26] to determine if other signaling components associated with PLCβ signaling are expressed in Type III taste cells. We found that the alpha G-protein Gαq which activates PLCβ is expressed in these cells. We also identified IP_3_R1 expression in Type III cells, which is a downstream target of PLC [26]. In Type II cells, PLCβ2 activates IP_3_R3 which causes a Ca^2+^ release to activate TRPM4 and TRPM5 (see Figure 1). Our search of the RNA-seq dataset did not find TRPM5 in Type III cells, but did identify TRPM4 expression, suggesting a possible downstream target of the Ca^2+^ release signal in the BR cells.

These data are surprising because it is currently thought that Type III cells primarily respond to sour or salt stimuli. Several studies focused on understanding Type III taste cells targeted and ablated PKD2L1 expressing cells since PKD2L1 is thought to be expressed in all Type III taste cells. They found that loss of these cells abolished the gustatory nerve responses to sour but did not alter gustatory nerve signaling to bitter, sweet and umami [22, 62]. A separate study reported that Type III cells are heterogeneous and do not uniformly express the same cell markers [29], so it may be that the BR cells are separate from the PKD2L1 expressing Type III cells. Recently, OTOP1, which is expressed in Type III taste cells, has been proposed as the sour receptor [54, 63, 64]. Nerve recordings from OTOP1-KO mice were significantly reduced for sour stimuli but also showed some reduction in their responsiveness to bitter, sweet and umami [64]. Taken together, these data highlight a heterogeneity in the characteristics of Type III taste cells that has not yet been well defined.

After taste receptors for bitter, sweet and umami were molecularly identified [65–69], multiple labs focused on identifying where these receptors are expressed in taste cells. While most taste receptors (T1Rs and T2Rs) are thought to be expressed only in Type II taste cells [4], there is some heterogeneity in their expression [70–74] and one RNA-seq dataset reported very low levels of expression for both T1R1 and T1R3 in some Type III cells [26]. This study evaluated 14 identified Type III cells, so it is possible that a larger analysis would identify more Type III cells that express these taste GPCRs [26]. In addition to these studies, multiple laboratories have independently concluded that the identified taste receptors are not solely responsible for transducing all bitter, sweet and umami stimuli. For instance, the detection of some carbohydrates is not impaired by the loss of T1R2 or T1R3, the identified sweet receptors [75–80], and umami stimuli appear to be transduced by receptors in addition to the T1R1+T1R3 heterodimer [81–89]. The glucose transporter which has been implicated in detecting sweet stimuli [90], was also expressed in some Type III cells in the RNA-seq profiling study. Our findings support the idea that there may be additional taste receptor mechanisms, outside of those canonically identified, involved in the detection of bitter, sweet and umami stimuli. While our study is not focused on identifying the receptors that activate the PLCβ3 signaling pathway, our data indicate that future studies are needed to identify which taste receptors are important in BR cells.

Even though the BR cells are only a subset of cells within the bud, our data suggest that these cells make a significant contribution to taste. We measured the neural activity in the NTS using c-Fos labeling as a marker for activated neurons to determine how the loss of PLCβ3 affects central processing of taste information from the periphery (Figure 6). We used oral infusions of taste stimuli in awake and behaving animals because it removes any potential confounds due to anesthetization that is used in other approaches. Since the NTS receives input from the gustatory neurons and is the first synaptic relay from the peripheral taste system, these data confirmed that loss of PLCβ3 in the taste cells significantly reduced the taste signal that is sent to the brain. The loss of PLCβ3 also caused very significant impairments in the behavioral responses to bitter, sweet and umami stimuli (Figure 7). These deficits were comparable to the loss of IP_3_R3 in the Type II cells which has a well-established role in the transduction of bitter, sweet and umami stimuli. The effects of losing either protein were specific to these taste stimuli and did not affect sodium taste (Figure 7E).

Since PLCβ3 and IP_3_R3 are not expressed in the NTS (Figure S6) or the geniculate ganglia [91], our data suggest that input from both of these signaling molecules in taste cells is required for normal taste behavior. Moreover, if Type II and BR taste cells independently signal to the central taste system, then the loss of input from either cell population would be predicted to reduce the NTS activity proportionally to their contribution in taste processing. However, our data do not support this prediction since the loss of either PLCβ3 (in BR cells) or IP_3_R3 (in Type II cells) reduced c-Fos labeling in the NTS to background (water) levels. This suggests that input from both Type II (IP_3_R3 expressing) *and* BR (PLCβ3 expressing) cells may be required for an optimal taste response. When either cell population is non-functional, there are severe deficits in both NTS activity and behavioral responses. This type of synergistic relationship is present in multiple sensory systems [92–98] but to our knowledge, it has not previously been described in the taste system. Future studies are needed to better understand this potential relationship.

The presence of broadly tuned taste cells raises an interesting question about how taste information is coded to the brain. One school of thought is that taste information is carried in a labeled line; individual taste cells respond to a particular taste stimulus and this specificity is maintained throughout the taste system [99]. If this is correct, then it is unclear what role BR cells would have in coding taste information. Conversely, another school of thought is that taste information is coded using an across fiber approach. Gilbertson et al [48] suggested that multicomponent messages have a greater capacity to transmit information. Encoding information with patterns of activity across broadly tuned cells allows for the transmission of more information than a system using specifically tuned cells [48, 100]. Having broadly tuned cells would allow the taste system to better discriminate between chemicals with similar characteristics. The presence of specialist and generalist cell populations within the taste system suggests that taste coding may be incorporating aspects of both labeled line and across fiber coding. Future studies are needed to address this possibility.

Our study has demonstrated that a subset of Type III cells responds to bitter, sweet and/or umami stimuli using a PLCβ3 signaling pathway. These cells are broadly tuned to multiple stimuli, including stimuli from distinct taste modalities. This subset of Type III cells appears to have an important role in taste since the loss of either IP_3_R3 (in Type II cells) or PLCβ3 (in BR cells) causes similar deficits in both NTS responses and taste driven behaviors. We conclude that peripheral taste transduction is more complex than is currently appreciated.

## EXPERIMENTAL PROCEDURES

### Mice

Animals were cared for in compliance with the University at Buffalo Institutional Animal Care and Use Committee. Multiple mouse lines were used in this study. Both sexes were used and mice ranged in age from 1 to 6 months. Taste cells for qPCR analyses were collected from C57BL/6 mice. The IP_3_R3-KO mouse was generated in a C57BL/6 background and was obtained from the Mutant Mouse Resources and Research Center (MMRRC:032884-JAX). This mouse has a targeted mutation in which exon 1 is replaced with a MAPT/GFP fusion [101] which results in the expression of GFP in place of a functional IP_3_R3 receptor. Thus, these mice lack a functional IP_3_R3 receptor and express GFP in cells that would otherwise have expressed IP_3_R3. Heterozygous mice express both GFP and the IP_3_R3 receptor. The PLCβ3-KO mouse was generously provided by Dr. Sang-Kyou Han [102]. The mutation in these mice disrupts the catalytic domain of phospholipase C and was generated in a 129SV agouti mouse strain that was crossed with CD1 mice [103]. These mice were maintained in this mixed background. Immunohistochemical analyses were also performed in the TRPM5-GFP mouse strain which expresses GFP in all taste cells that express TRPM5. These mice have been backcrossed into the C57BL/6 background and were used as another marker of Type II taste cells [15] to evaluate the expression patterns of PLCβ3. These mice were generously provided by Dr. Robert Margolskee.

### Taste Receptor Cell Isolation

Taste receptor cells were harvested from CV, Fol and Fun papillae of adult mice as previously described [23, 104–109]. Briefly, mice were sacrificed using carbon dioxide and cervical dislocation. Tongues were removed and an enzyme solution containing 0.7mg/mL Collagenase B (Roche, Basel, Switzerland), 3mg/mL Dispase II (Roche), and 1mg/mL Trypsin Inhibitor (Sigma-Aldrich, St. Louis, MO) was injected beneath the lingual epithelium. After the tongues were incubated in oxygenated Tyrode’s solution for approximately 17 min, the epithelium was peeled from the underlying muscle and pinned serosal side up before it was incubated in Ca^2+^-free Tyrode’s for approximately 25 min. Cells were removed from taste papillae using capillary pipettes with gentle suction and placed onto coverslips coated with Cell-Tak (Corning, Corning, NY).

### Ca^2+^ Imaging

All measurements of intracellular calcium (Ca^2+^) were performed in isolated taste receptor cells that were no longer in contact with other taste cells. Cells were loaded for 20 minutes at room temperature (RT) with 2 μM Fura2-AM (Molecular Probes, Invitrogen, Carlsbad, CA) containing Pluronic F-127 (Molecular Probes). Loaded cells were then washed in Tyrode’s solution under constant perfusion for 20min. Multiple taste stimuli and high potassium (50mM KCl) solutions were individually applied and Ca^2+^ responses were recorded. Cells were visualized using an Olympus IX73 microscope with a 40x oil immersion lens and images were captured with a Hamamatsu ORCA-03G camera (Hamamatsu Photonics K.K., SZK Japan). Excitation wavelengths of 340nm and 380nm were used with an emission wavelength of 510nm. Cells were kept under constant perfusion using a gravity flow perfusion system (Automate Scientific, San Francisco, CA). Images were collected every 4s using Imaging Workbench 6.0 (Indec Biosystems, Santa Clara, CA). Experiments were graphed and analyzed using Origin 9.2 software (OriginLab, Northhampton, MA).

Intracellular Ca^2+^ levels were measured as a ratio of fluorescence intensities. Fluorescence values were calibrated using the Fura-2 Ca^2+^ Imaging Calibration kit (Invitrogen). The effective dissociation constant K_d_ was calculated to be 180nM, which was used in the following equation to calculate Ca^2+^ concentration:

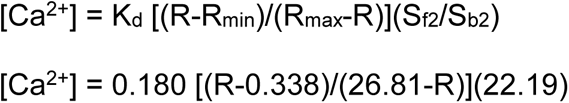

R is the ratio value of fluorescence obtained after exciting cells at 340 and 380nm. Data from cells were analyzed if the cell had a stable Ca^2+^ baseline within the range of 65nM and 200nM. An evoked response was defined as measurable if the increase in fluorescence was at least two standard deviations above baseline.

### Immunohistochemistry

Mice were deeply anesthetized by intraperitoneal injections of sodium pentobarbitol, 40 mg/kg (Patterson Veterinary, Mason, MI). Mice were then injected intracardially with heparin (Sigma) and 1% sodium nitrate (Sigma) followed by perfusion with approximately 30mL of 4% paraformaldehyde (Electron Microscopy Sciences, Ft. Washington, PA) in 0.1M phosphate buffer (PB), pH 7.2. After perfusion, the tongues were removed and placed in 4% paraformaldehyde/0.1M PB for 1-2h followed by a 4°C overnight incubation in 20% sucrose/0.1M PB, pH 7.2. For some experiments, tongues were immersion fixed overnight in 4% paraformaldehyde/0.1M PB, pH 7.2 at 4°C with 20% sucrose. Regardless of the fixation method, the next day, 40µm sections were cut and washed in PBS 3X10 min at RT. For some experiments, antigen retrieval was performed by placing sections in 10 mM sodium citrate, pH 8.5 at 80°C for 5 min. This was done to disrupt the cross-bridges formed by fixation and expose antigen binding sites.

Sections were incubated in blocking solution (0.3% Triton X-100, 1% normal goat serum and 1% bovine serum albumin in 0.1M PB) for 1-2h at RT. Primary antibody was added to the sections in blocking solution and incubated for 2 hours at RT followed by overnight exposure to primary antibody at 4°C. Controls with no primary antibody were included in each experiment. All primary antibodies were diluted in blocking solution. Mouse anti-IP_3_R3 (Transduction Labs, Lexington, KY) was used at 1:50 following antigen retrieval. Rabbit anti-PLCβ3 (Abcam) was used at 1:200, rabbit anti-PLCβ2 (Santa Cruz Laboratories, Santa Cruz, CA) was used at 1:1000, rabbit anti-gustducin (Santa Cruz) was used at 1:200 and anti-NTPDase2 (1:100[110]. Mouse anti-SNAP-25 (Genway Biotech, San Diego, CA) was used at 1:200 following antigen retrieval. Following overnight incubation in primary antibody, sections were washed in PBS 3X10 min at RT and then incubated with the appropriate secondary antibody (Cy5, 1:500; Rhod, 1:250; Jackson ImmunoResearch Laboratories, West Grove, PA) at RT for 2h in the dark. Controls were performed for double labeling experiments to ensure secondary antibodies were not binding to primary antibodies raised in different organisms. After secondary antibody incubation, sections were washed in PBS (3×10 min) and mounted on Superfrost Plus slides (VWR, Radnor, PA) using Fluoromount G (Southern Biotechnology Associates, Birmingham, AL) and coverslipped. All images were obtained using a Zeiss LSM 710 Confocal Microscope (Zeiss, Oberkochen, Germany). Stacks were collected using Zen software (Zeiss) and images were processed using Adobe Photoshop CS5 software adjusting only brightness and contrast.

### Real-Time PCR of Isolated Taste Cells

Taste receptor cells from CV, Fol or Fun papillae were isolated from the papillae as described above and then centrifuged for 20min at 13,000 RPM. RNA was purified using the NucleoSpin RNA XS Kit (Macherey-Nagel, Düren, Germany) according to kit instructions. PCR analysis was performed for GAPDH to ensure sample quality and check for genomic contamination. Contaminated samples were discarded and new samples were collected. Real-Time PCR was performed using a BioRad MiniOpticon system (Bio-Rad Laboratories, Hercules, CA), with BioRad SYBR Green reagents (Bio-Rad Laboratories). Primers used for these experiments were: **PLCβ2**, *Fwd*: CAATTGAGGGGCAGCTGAGA *Rev*: TTCTAGGCTGCATCTGGGC; **PLCβ3**, *Fwd*: TCCTGGTGGTCAGGGAT *Rev*: CTGCCTGTCTCTGCTATCCG; **GAPDH** *Fwd*: ACAGTCAGCCGCATCTTCTT, *Rev*: ACGACCAAATCCGTTGACTC.

For real-time PCR analyses, each sample was run in triplicate. If there was more than 5% difference between the replicates, the data were discarded. Multiple biological repeats were used for each papillae type (CV, n=6; Fol, n=6; Fun, n=5). Data was normalized to GAPDH expression for each sample to correct for any loading differences and reported as fold differences.

### Oral Infusions

Wild type, IP_3_R3-KO and PLCβ3-KO mice (n=3 for each) were tested to measure the effects of quinine stimulation on the c-Fos immunoreactivity in the nucleus of the solitary tract (NTS). Water infusions (n=3) were performed to measure background responses. Surgical procedures were conducted using sterile technique while mice were anesthetized via 2% - 4% isoflurane anesthesia. Guided by a 21G needle, a small length of PE-90 tubing was passed through the animal’s cheek, emerging into the oral cavity by the rear molars. The tubing was threaded through Teflon washers and heat flared to secure on both the external and oral sides. Mice were given an analgesic (0.5mg/ml of carprofen) and allowed to recover. After recovery, they were infused with either quinine (5mM) or water into the oral catheter for 30 minutes (0.2ml/min, 30 min infusion). Following the infusion, animals were returned to their cages and left undisturbed for 45 minutes.

### Brain Histology and Analysis

After the 45 minute post-infusion period, mice were perfused as described above and the hindbrains were removed. The following day, hindbrains were sectioned into 40μm coronal sections which were washed in TBS (pH 7.5), 3×10 min each. Sections were incubated in 1:100 H_2_O_2_ in TBS for 15 min, followed by 3X10 min TBS washes. Sections were then incubated in anti-c-Fos (ab190289; Abcam, 1:1000) diluted in blocking buffer (3% Normal Donkey Serum in TBS-TritonX) for 1 hr at RT. This was followed by an overnight incubation at 4°C. The next day, sections were washed 3×10 min in TBS and then incubated for 2 hrs in anti-rabbit secondary antibody (711-065-152, Jackson Immunoresearch, 1:1000) in blocking buffer. After incubation, sections were kept in an avidin-biotin mixture (Elite kit; Vector Laboratories) in TBS-TritonX for 1 hr. Tissue sections were washed (3×10 min in TBS) and then stained with DAB (3,3’-diaminobenzidine-HCL; Vector Laboratories) for 5 min. Stained tissue sections were washed (3×5 min in TBS) and mounted onto slides using Fluoromount-G (Southern Biotechnology Associates). Sections were examined using a light microscope (10-100 X) equipped with a digital camera.

Analyses of digital images were performed on the intermediate rostral nucleus tractus solitarius (iRNTS; ∼500 µm caudal to the dorsal cochlear nucleus) which receives afferents from both the glossopharyngeal and chorda tympani nerves and displays dense c-Fos-like immunoreactivity in response to intraoral delivery of quinine [35, 111]. A template tracing of the nucleus was made by an experimenter blind to genotype and stimulus condition and this template was applied to all sections. The number of FLI-positive neurons within the template was counted by hand by an experimenter blind to genotype. Data were analyzed with a one way ANOVA with follow up Student’s t tests to identify individual differences. Significance level was set at p<0.05.

### Analysis of Licking Behavior

Unconditioned licking responses to varying concentrations of taste stimuli were recorded in a test chamber designed to measure brief-access licking (Davis MS80 Rig; Dilog Instruments and Systems, Tallahassee, FL). This apparatus consisted of a Plexiglas cage with a wire mesh floor. An opening at the front of the cage allowed access to one of sixteen spill-proof glass drinking tubes that reside on a sliding platform. A mechanical shutter opened and closed to allow the mouse access to one of the tubes for a user-specified length of time. A computer controlled the movement of the platform, order of tube presentation, opening and closing of the shutter, duration of tube access and interval between tube presentations. Each individual lick was detected by a contact lickometer and recorded on a computer via DavisPro collection software (Dilog Instruments and Systems).

Mice were adapted to the test chamber and trained to drink from the sipper tubes for 5 consecutive days as previously described [112, 113]. During training, mice were 20-h water deprived. On the first day of training, the mouse was presented with a single stationary bottle of water for 30 min. On the second day, a tube containing water was presented but this time the mouse was given 180s to initiate licking and once licking was recorded the mouse was given 30s access to the tube. At the conclusion of either the 30s access or the 180s limit, the shutter was closed again for 10s. Each of the 8 tubes, all containing water, was presented 3 times. During the remaining three days of training, the mouse was given 30 min to initiate licking to one of eight tubes of water. Once the mouse began licking, it was given 10s to lick before the shutter closed for 10s, after which a new tube was presented.

During testing, animals were allowed to take as many trials as possible in 30 min. Mice were tested on varying concentrations of sucrose (0,3,10,30,60,100,300,1000 mM), acesulfame K (0,0.5,2,6,8,16,20,32 mM), MSG with 10 µM amiloride (0,25,50,75,100,400,600,800 mM), denatonium benzoate (0,0.1,0.3,1,3,5,10,20 mM), and NaCl (0,3,10,30,60,100,300,1000 mM), in that order. Each stimulus was presented in randomized blocks on Monday, Wednesday and Friday in a single week. Animals were 22-h water deprived for all testing except sucrose, when animals were tested water replete. Once the animal began licking the tube, they were allowed 10 seconds of access before the shutter closed.

For stimuli tested in the water deprived condition (acesulfame K, MSG + amiloride, denatonium benzoate, and NaCl), lick ratios were calculated by dividing the average number of licks at each concentration by the average number of licks to water. For stimuli tested while the animals were water replete (sucrose) licks scores were calculated by subtracting the average number of licks at each concentration by the average number of licks to water. These corrections are used to standardize for individual differences in lick rate and are based on water need. Lick scores and licks relative to water are compared by repeated measures ANOVA with genotype as between factors variable and concentration as a repeated measures within factors variable. Significant interaction terms were followed by Tukey’s Honestly Significant Difference tests. Statistical analyses were performed in Statistica.

### Solutions

All chemicals were purchased from Sigma Chemical (Sigma-Aldrich, St. Louis, MO) unless otherwise noted. Tyrode’s solution contained 140mM NaCl, 5mM KCl, 3mM CaCl_2_, 1mM MgCl_2_, 10mM HEPES, 10mM glucose, and 1mM pyruvate, pH 7.4. Ca^2+^-free Tyrode’s contained 140mM NaCl, 5mM KCl, 2.7mM BAPTA, 2mM EGTA, 10mM HEPES, 10mM glucose, 1mM pyruvate, pH 7.4. Nominal Ca^2+-^free Tyrode’s contained 140mM NaCl, 5mM KCl, 10mM HEPES, 10mM glucose, 1mM pyruvate, pH 7.4. Hi KCl solution contained 50mM KCl, 90mM NaCl, 3mM CaCl_2_, 1mM MgCl_2_, 10mM HEPES, 10mM glucose, 1mM pyruvate, pH 7.4. All taste-solutions, 2µM U73122 (Tocris, Bristol, United Kingdom), and 2µM thapsigargin (Tocris) were prepared in normal Tyrode’s. Multiple taste stimuli were analyzed for bitter (5mM denatonium benzoate (Den), sweet (20mM sucralose, 2mM saccharin, 50mM sucrose, 20mM Acesulfame K (Ace K)), and umami (10mM monopotassium glutamate (MPG)). 50mM sucrose contained 90mM NaCl instead of 140mM. Salt (250mM sodium chloride, NaCl) and sour (50mM citric acid, CA, pH4) stimuli were tested using the same protocol as Lewandowski et al [18].

### Statistics

Analysis of the taste responsiveness was performed using an interactive Chi-square analysis with Yate’s correction for continuity [114]. Significant differences were reported if p<0.05. For real-time PCR analyses, a one-way ANOVA with p<0.05 set as the limit of significance was used to identify any significant differences between the relative expression of PLCβ2 and PLCβ3 in the different papillae types. Comparisons between two experimental conditions were made using a two-tailed Student’s T test with p<0.05 set as the limit of significance.

### Colocalization analysis

Image stacks (n=5, 1 µm each) were acquired from labelled sections obtained from three different animals and regions of interest (ROI) were drawn to include the area inside clearly identifiable taste buds. Colocalization analysis was performed on the ROIs using ImageJ Fiji software (NIH). Colocalization was determined based on Pearson’s coefficient using the Colocalization_Finder plugin. If the Pearson’s coefficient value was greater than 0.9, we considered it to be 100% colocalization due to the variability in immunofluorescence intensity. If the Pearson’s coefficient was less than 0.05, we considered this to be no colocalization. A Pearson’s coefficient higher than 0.05 but lower than 0.9 was considered as partial colocalization.

## ACKNOWLEDGMENTS

The authors thank Drs. Sue Kinnamon and Stefan Roberts for comments. This work was supported by NSF1256950 to KFM. The authors thank Alan Siegel and the UB North Campus Imaging facility funded by NSF-MRI DBI0923133 for the confocal images. The authors thank Jhanna Flora for generating our model images.

## AUTHOR CONTRIBUTIONS

DDB and EDB did live cell imaging, immunohistochemistry, data analysis and contributed to writing the manuscript. DDB also did infusion experiments. LEM performed brief-access lick behavior experiments, infusion experiments and data analysis. KEK performed infusion experiments and data analysis. GCL conceived and analyzed infusion experiments. ARN did immunohistochemistry, qPCR and data analysis. ZCA and BRK did live cell imaging and data analysis. BTK did immunohistochemistry. AMT supervised brief-access lick behavior, infusion experiments, analyzed and interpreted the data, and edited the manuscript. KFM conceived and supervised the project, did live cell imaging, data analysis and wrote the manuscript.

The authors declare no competing interests.

---

**Figure S1.**
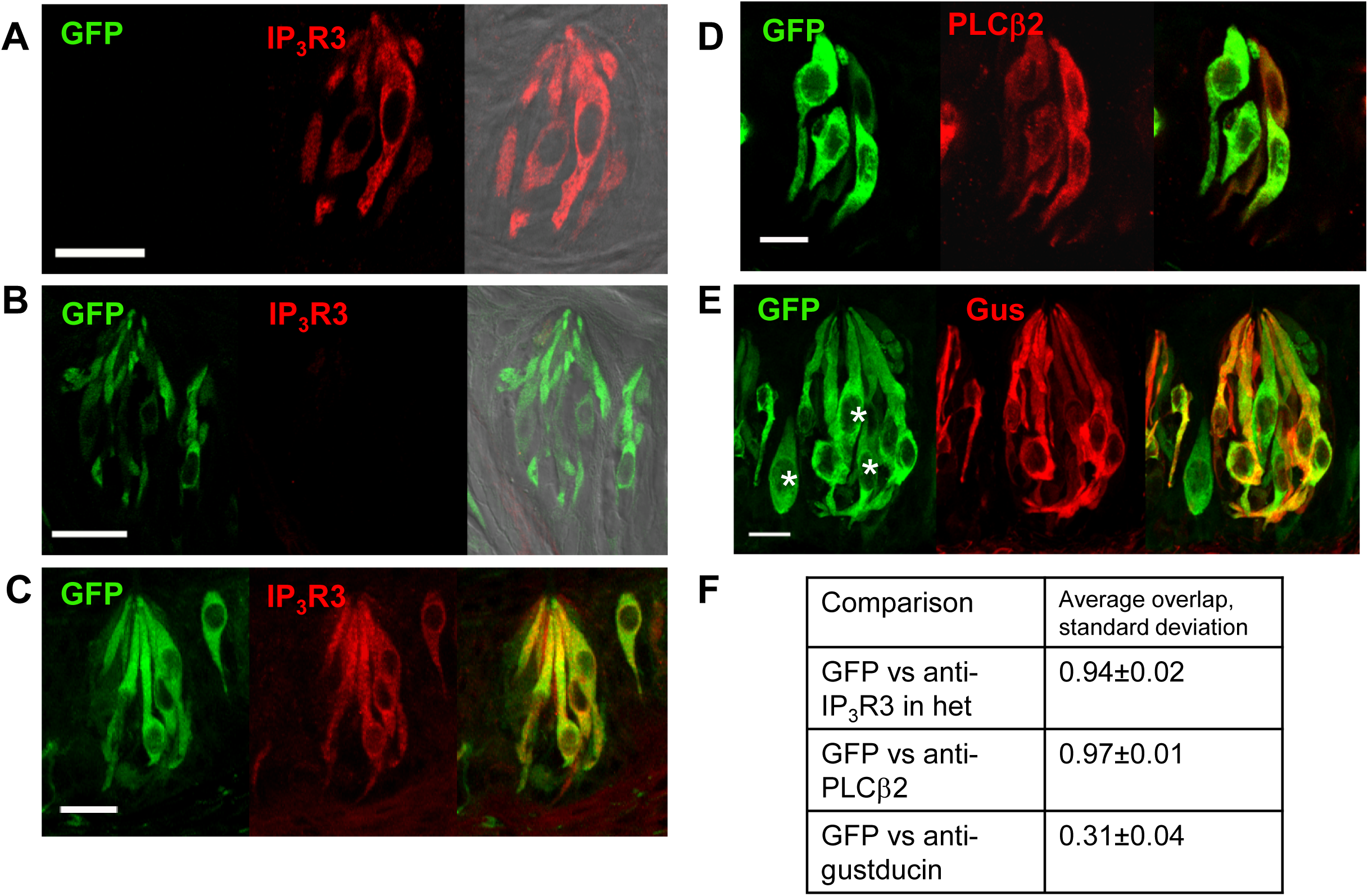
Characterization of the IP_3_R3-KO mice. A) Laser scanning confocal micrographs (LSCMs, stack of 5 slices, 1µm each) from WT mice identified anti-IP_3_R3 labeling in taste receptor cells from the CV (n=3). B) IP_3_R3-KO mice express GFP in lieu of IP_3_R3 and were not labeled by anti-IP_3_R3 (n=6). C) LSCMs of the IP_3_R3-het mouse identified strong co-localization between anti-IP_3_R3 labeling and GFP expression. D) Anti-PLCβ2 labeling in IP_3_R3-KO mice found that PLCβ2 co-localizes with GFP, indicating that IP_3_R3-KO-GFP is specific to Type II cells (LSCMs: stack of 5 slices, 1µm each; n=4). E) α-gustducin is present in a subset of the IP_3_R3-KO-GFP taste cells in the CV (LSCMs: stack of 10 slices, 1µm each; n=3). Asterisks identify some GFP expressing cells that do not express gustducin. Scale bars = 20 µm. F) Co-localization analysis identified the average (± standard deviation) overlapping expression for each target protein with the GFP expression, n=3 for each.

**Figure S2.**
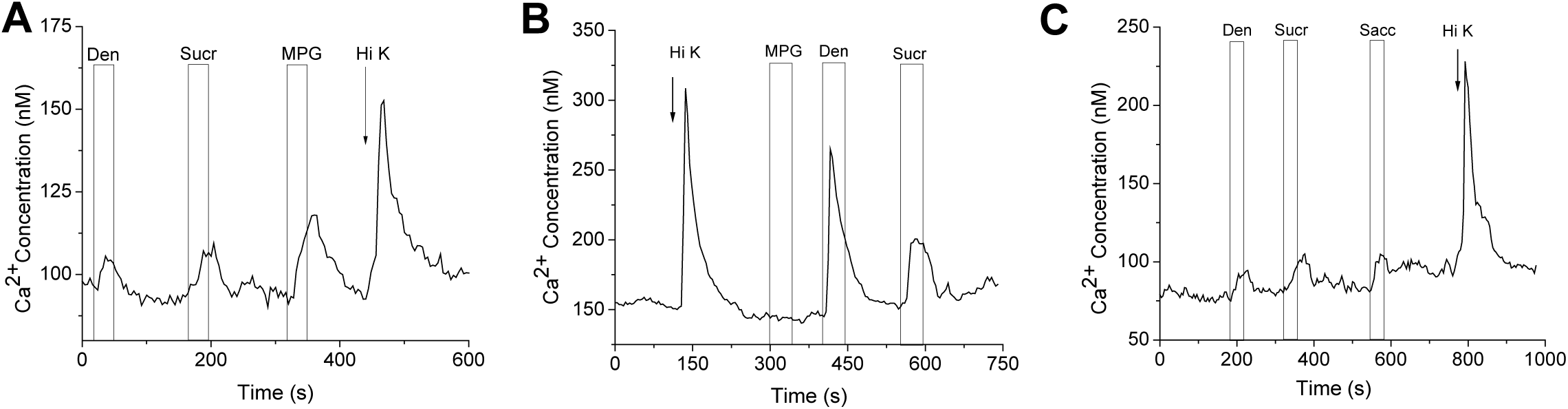
Dual responsive cells are present in multiple mouse lines. Control imaging experiments were performed using GAD67-GFP mice and C57BL/6 mice. GAD67-GFP is expressed in a large subset of Type III mouse taste cells [27]. A-B) Representative traces of BR taste cells that responded to bitter (denatonium=Den), sweet (sucralose=Sucr) and/or umami stimuli (MPG) and 50mM KCl (Hi K) in GAD67-GFP mice. BR taste cells were present in both GAD67-GFP + (A) and GFP-(B) taste cells. C) Experiments in C57BL/6 mice also identified the presence of BR taste cells.

**Figure S3.**
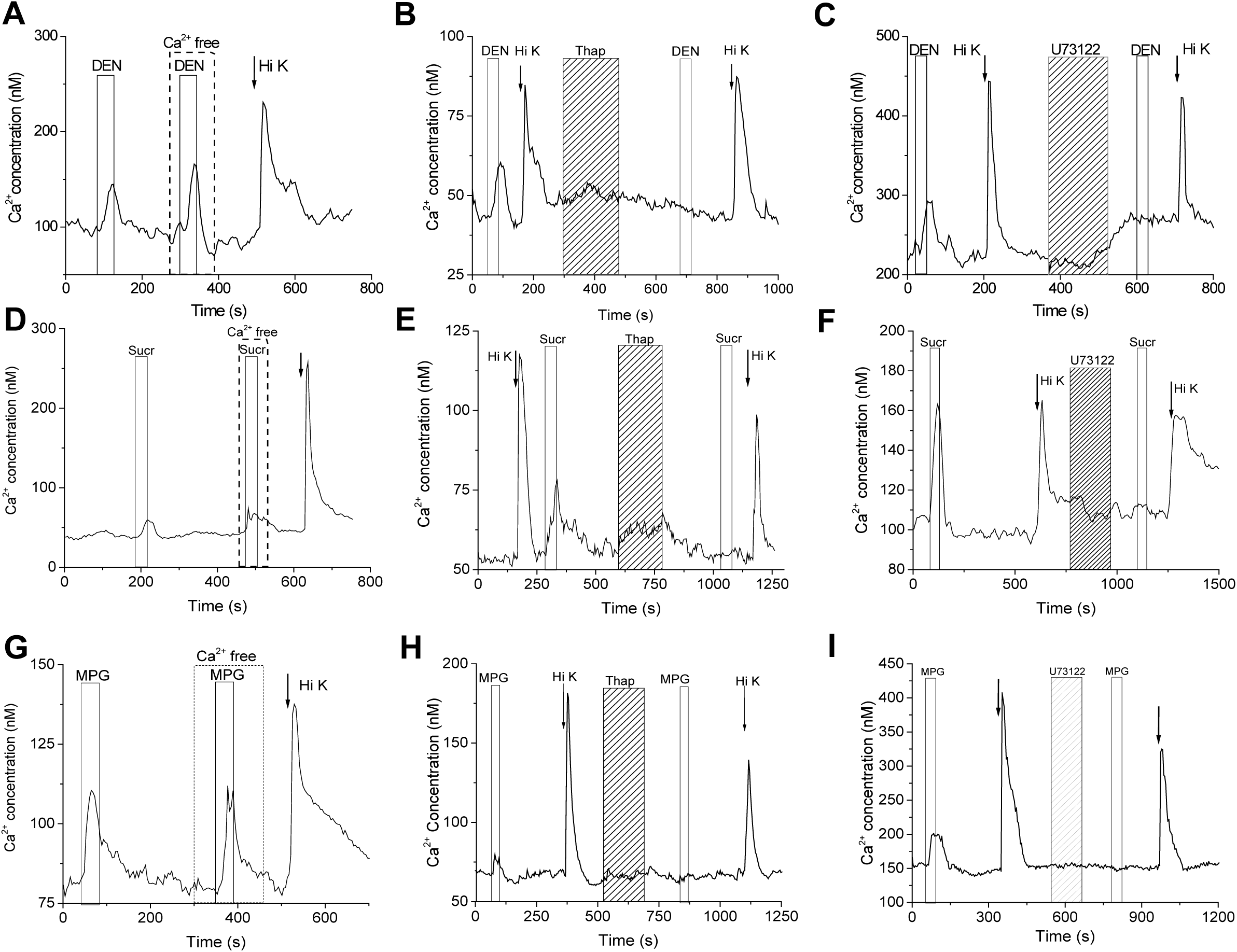
Taste-evoked Ca^2+^ release in IP_3_R3-KO mice is dependent upon PLC activity and Ca^2+^ release from internal stores. Representative data related to Figure 4. Open columns represent the time that the taste stimulus is presented. The application of Ca^2+^ free Tyrode’s is indicated by the dashed lines. The stimulus presented during this time is also in Ca^2+^ free Tyrode’s. The gray hatched columns represent the application of either thapsigargin (Thap) or U73122, both of which are irreversible inhibitors. A) Bitter-evoked taste responses (5mM Den) persist in the absence of extracellular calcium (Ca^2+^-free) and are abolished by the SERCA pump inhibitor thapsigargin (B) as well as the PLC blocker U73122 (C). D) Responses to sweet stimuli (20mM sucralose, Sucr) persist in Ca^2+^-free and are abolished by thapsigargin (E) and U73122 (F). G) Umami stimuli (10mM MPG) persist in Ca^2+^-free and were abolished by thapsigargin (H) and U73122 (I).

**Figure S4.**
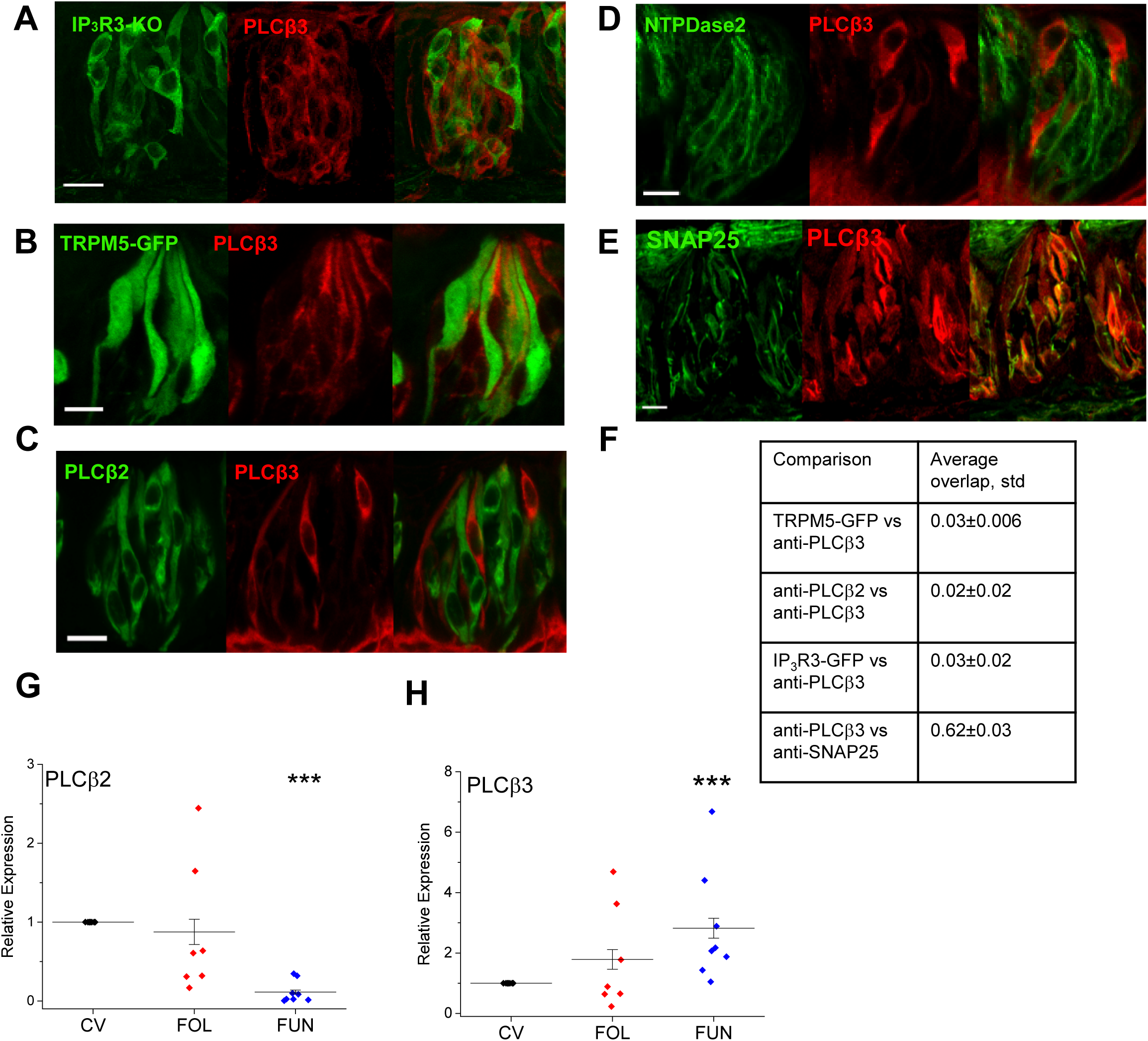
Expression of PLCβ3 in taste cells. A) Laser scanning confocal micrographs (LSCMs, stack of 5 slices, 1µm each) of PLCβ3 immunostaining in the IP_3_R3-KO-GFP mice reveal that PLCβ3 is expressed in a separate population from the GFP positive taste cells in the CV. B) Anti-PLCβ3 labeling in the CV of TRPM5-GFP mice determined that PLCβ3 is expressed in taste cells lacking GFP expression (LSCMs: stack of 5 slices, 1µm each; n=4). C) Co-labeling with anti-PLCβ2 and anti-PLCβ3 in the CV of C57BL/6 mice revealed that these PLCβs are expressed in separate taste cell populations (LSCMs: stack of 5 slices, 1µm each; n=3). D) Co-labeling with anti-NTDPase2 and anti-PLCβ3 in the CV of C57BL/6 mice determined that these markers are expressed in separate taste cell populations (LSCMs: stack of 5 slices, 1µm each; n=3). Scale bar = 20µm. E) Immunohistochemical analyses (LSCMs: stack of 5 slices, 1µm each) using anti-PLCβ3 and anti-SNAP25 revealed some co-localization between PLCβ3 and SNAP25 in CV papillae. Scale bar = 10µm. F) Co-localization analysis identified the average (± standard deviation) overlapping expression for PLCβ3 with TRPM5-GFP, anti-PLCβ2, IP_3_R3-GFP or anti-SNAP25 expression, n=3 for each. mRNA was isolated from taste cells originating in the different papillae types from C57BL/6 mice. Taste cells were analyzed from at least five different mice for each. Values were normalized to GAPDH expression and are presented as a ratio to values from the CV papillae for (G) PLCβ2 and (H) PLCβ3. (***, p<0.001).

**Figure S5.**
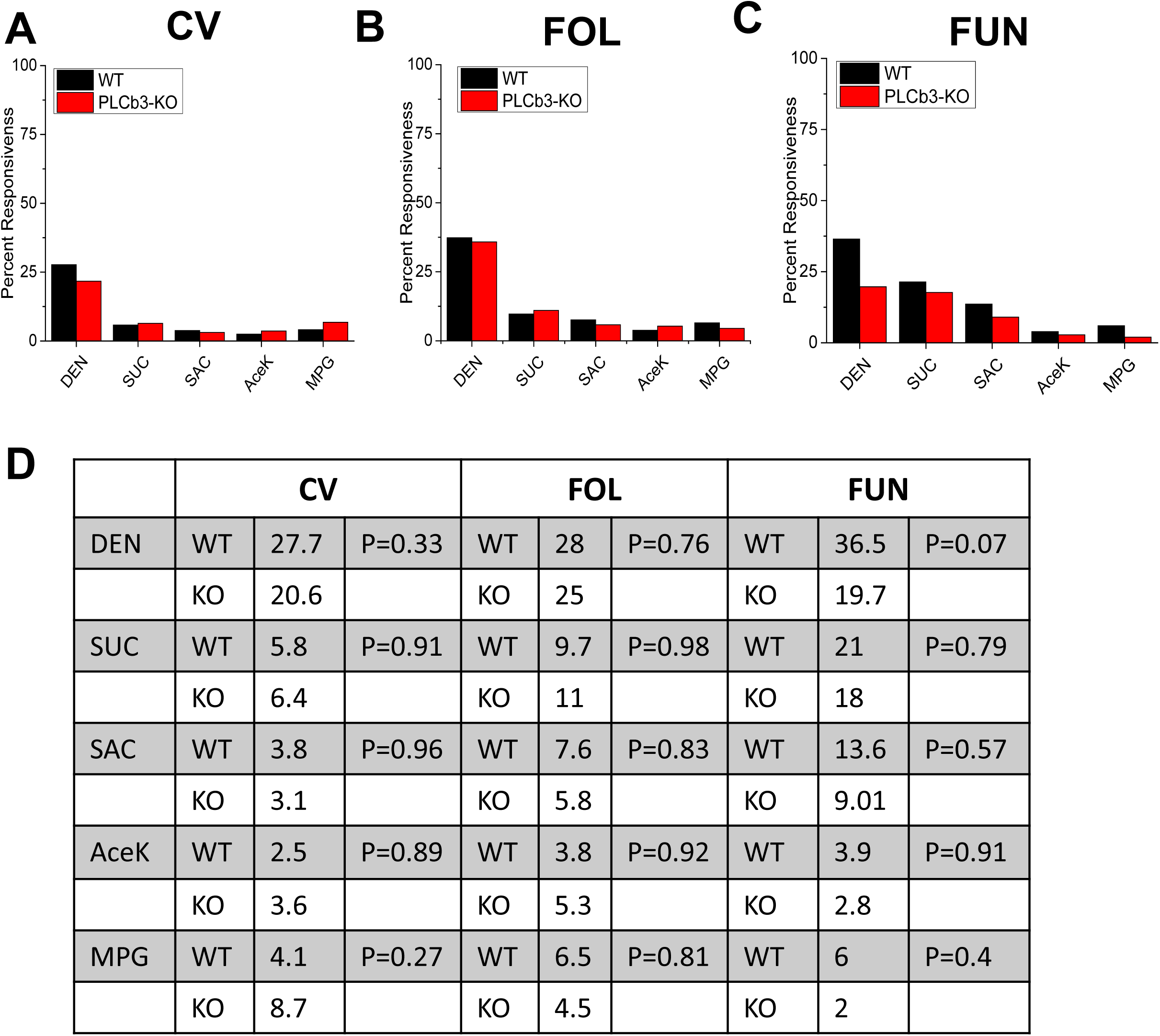
Loss of PLCβ3 expression does not affect Type II TRC responses. Chi square analysis with Yate’s correction for continuity was used to compare the frequency of evoked Ca^2+^ responses to different taste stimuli between wild type (black bars) and PLCβ3-KO (red bars) mice for taste cells from CV (A), Fol (B) and Fun (C) papillae. D). Table of the stimulus response frequency values for each papillae type in WT and KO mice. P values for each comparison are also shown. No significant differences were found for any of the comparisons.

**Figure S6.**
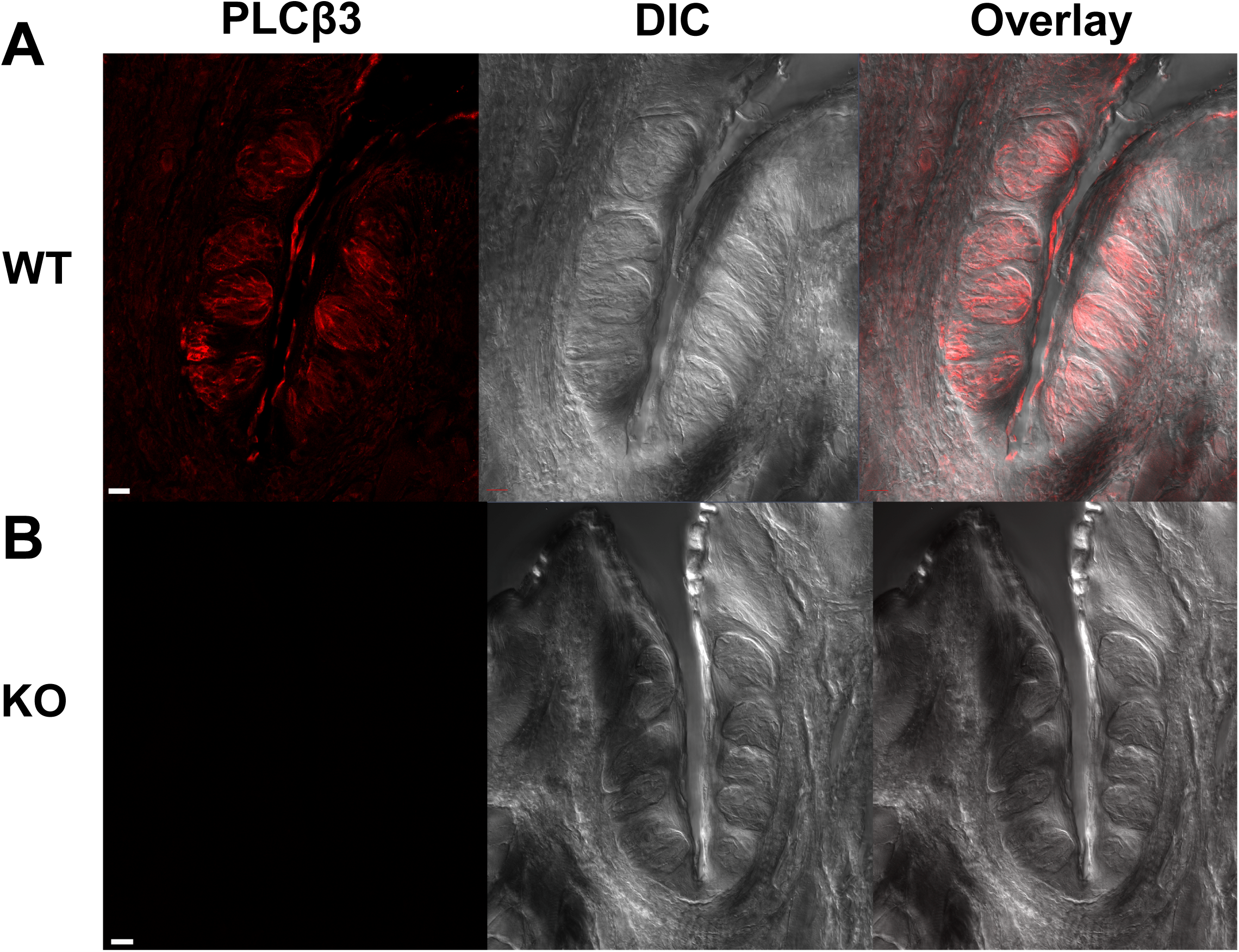
Characterization of the PLCβ3-KO mice. A) LSCMs (stack of 5 slices, 1µm each) from WT mice identified anti-PLCβ3 labeling in taste receptor cells from the CV (n=3). B) CV taste cells from the PLCβ3-KO mice were not labeled by anti-PLCβ3 (n=3). Scale bar = 10µm.

**Figure S7.**
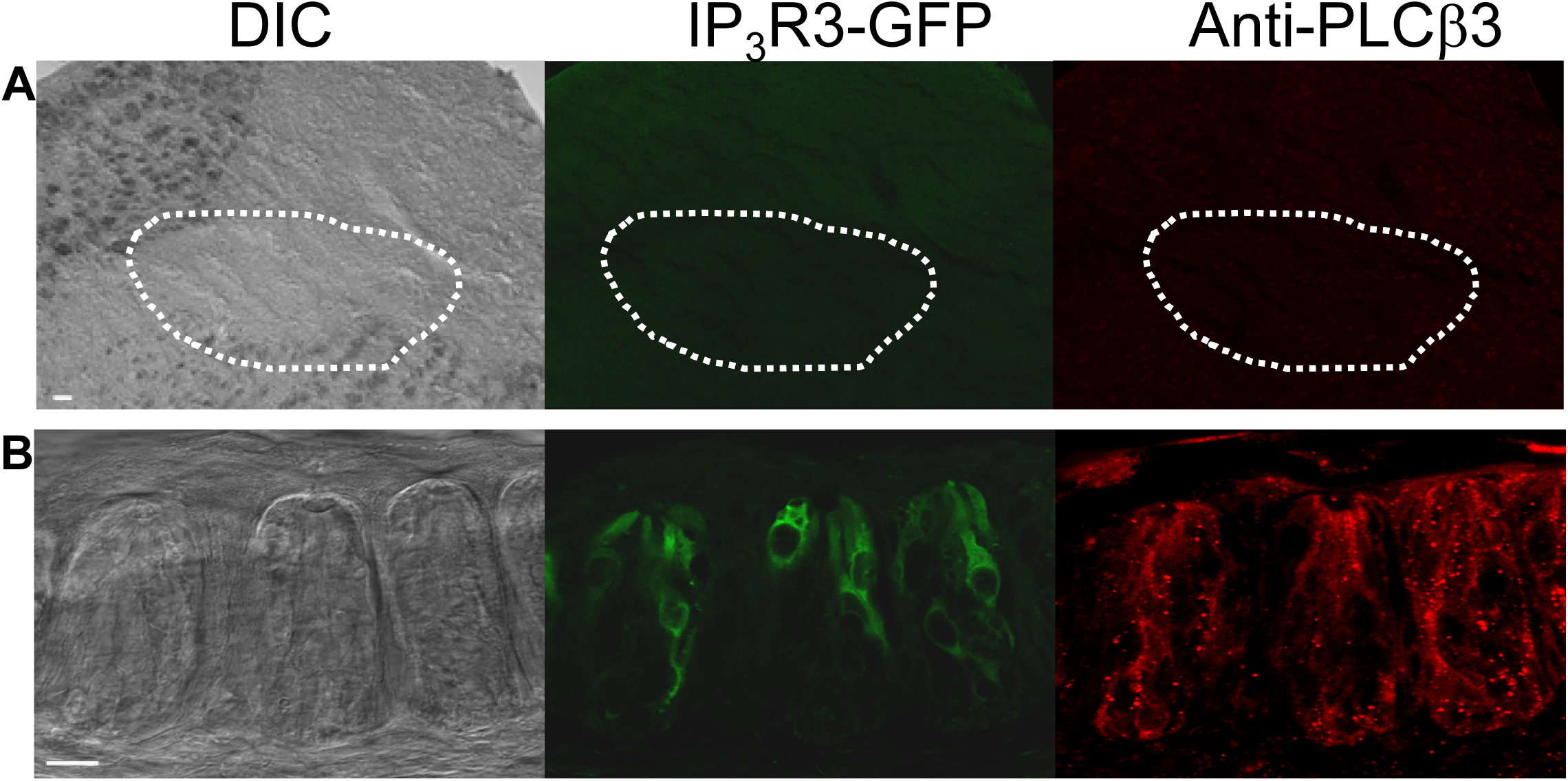
Neither IP_3_R3 nor PLCβ3 is expressed in IRNTS. CV papillae and IRNTS sections were evaluated in parallel. A) Analyses of the brain sections from the IP_3_R3-KO mice revealed no GFP labelling in the IRNTS. Immunohistochemical analysis using anti-PLCβ3 also did not detect any PLCβ3 expression. Scale bar=50µm. B) LSCM analyses (n=5 sections, 1 µm each) of the tongues from the same mice identified the expression of IP_3_R3-KO-GFP and PLCβ3 in the taste receptor cells from the CV papillae (n=3). Scale bar=10µm.

## REFERENCES

1. Finger TE. Cell types and lineages in taste buds. Chem Senses. 2005;30 Suppl 1:i54–5. doi: 10.1093/chemse/bjh110. PubMed PMID: 15738192.

2. Lawton DM, Furness DN, Lindemann B, Hackney CM. Localization of the glutamate-aspartate transporter, GLAST, in rat taste buds. Eur J Neurosci. 2000;12(9):3163–71. PubMed PMID: 10998100.

3. Miyoshi MA, Abe K, Emori Y. IP(3) receptor type 3 and PLCbeta2 are co-expressed with taste receptors T1R and T2R in rat taste bud cells. Chem Senses. 2001;26(3):259–65. PubMed PMID: 11287386.

4. Zhang Y, Hoon MA, Chandrashekar J, Mueller KL, Cook B, Wu D, et al. Coding of sweet, bitter, and umami tastes: different receptor cells sharing similar signaling pathways. Cell. 2003;112(3):293–301. PubMed PMID: 12581520.

5. Yan W, Sunavala G, Rosenzweig S, Dasso M, Brand JG, Spielman AI. Bitter taste transduced by PLC-beta(2)-dependent rise in IP(3) and alpha-gustducin-dependent fall in cyclic nucleotides. Am J Physiol Cell Physiol. 2001;280(4):C742–51. PubMed PMID: 11245589.

6. Dutta Banik D, Martin LE, Freichel M, Torregrossa AM, Medler KF. TRPM4 and TRPM5 are both required for normal signaling in taste receptor cells. Proc Natl Acad Sci U S A. 2018;115(4):E772–E81. doi: 10.1073/pnas.1718802115. PubMed PMID: 29311301.

7. Taruno A, Vingtdeux V, Ohmoto M, Ma Z, Dvoryanchikov G, Li A, et al. CALHM1 ion channel mediates purinergic neurotransmission of sweet, bitter and umami tastes. Nature. 2013;495(7440):223–6. doi: 10.1038/nature11906. PubMed PMID: 23467090; PubMed Central PMCID: PMC3600154.

8. Liu D, Liman ER. Intracellular Ca2+ and the phospholipid PIP2 regulate the taste transduction ion channel TRPM5. Proc Natl Acad Sci U S A. 2003;100(25):15160–5. PubMed PMID: 14657398.

9. Zhang Z, Zhao Z, Margolskee R, Liman E. The transduction channel TRPM5 is gated by intracellular calcium in taste cells. J Neurosci. 2007;27(21):5777–86. Epub 2007/05/25. doi: 27/21/5777 [pii] 10.1523/JNEUROSCI.4973-06.2007. PubMed PMID: 17522321.

10. Perez CA, Huang L, Rong M, Kozak JA, Preuss AK, Zhang H, et al. A transient receptor potential channel expressed in taste receptor cells. Nat Neurosci. 2002;5(11):1169–76. PubMed PMID: 12368808.

11. Clapp TR, Yang R, Stoick CL, Kinnamon SC, Kinnamon JC. Morphologic characterization of rat taste receptor cells that express components of the phospholipase C signaling pathway. J Comp Neurol. 2004;468(3):311–21. PubMed PMID: 14681927.

12. Yee CL, Yang R, Bottger B, Finger TE, Kinnamon JC. “Type III” cells of rat taste buds: immunohistochemical and ultrastructural studies of neuron-specific enolase, protein gene product 9.5, and serotonin. J Comp Neurol. 2001;440(1):97–108. Epub 2001/12/18. doi: 10.1002/cne.1372 [pii]. PubMed PMID: 11745610.

13. Simon SA, de Araujo IE, Gutierrez R, Nicolelis MA. The neural mechanisms of gustation: a distributed processing code. Nat Rev Neurosci. 2006;7(11):890–901. PubMed PMID: 17053812.

14. Roper SD. Signal transduction and information processing in mammalian taste buds. Pflugers Arch. 2007;454(5):759–76. PubMed PMID: 17468883.

15. Clapp TR, Medler KF, Damak S, Margolskee RF, Kinnamon SC. Mouse taste cells with G protein-coupled taste receptors lack voltage-gated calcium channels and SNAP-25. BMC Biol. 2006;4:7. PubMed PMID: 16573824.

16. Chang RB, Waters H, Liman ER. A proton current drives action potentials in genetically identified sour taste cells. Proc Natl Acad Sci U S A. 2010;107(51):22320–5. doi: 10.1073/pnas.1013664107. PubMed PMID: 21098668; PubMed Central PMCID: PMCPMC3009759.

17. Ye W, Chang RB, Bushman JD, Tu YH, Mulhall EM, Wilson CE, et al. The K+ channel KIR2.1 functions in tandem with proton influx to mediate sour taste transduction. Proc Natl Acad Sci U S A. 2016;113(2):E229–38. doi: 10.1073/pnas.1514282112. PubMed PMID: 26627720; PubMed Central PMCID: PMCPMC4720319.

18. Lewandowski BC, Sukumaran SK, Margolskee RF, Bachmanov AA. Amiloride-Insensitive Salt Taste Is Mediated by Two Populations of Type III Taste Cells with Distinct Transduction Mechanisms. J Neurosci. 2016;36(6):1942–53. doi: 10.1523/JNEUROSCI.2947-15.2016. PubMed PMID: 26865617; PubMed Central PMCID: PMCPMC4748077.

19. Huang YA, Maruyama Y, Stimac R, Roper SD. Presynaptic (Type III) cells in mouse taste buds sense sour (acid) taste. J Physiol. 2008;586(12):2903–12. doi: 10.1113/jphysiol.2008.151233. PubMed PMID: 18420705; PubMed Central PMCID: PMCPMC2517205.

20. Huang AL, Chen X, Hoon MA, Chandrashekar J, Guo W, Trankner D, et al. The cells and logic for mammalian sour taste detection. Nature. 2006;442(7105):934–8. doi: 10.1038/nature05084. PubMed PMID: 16929298; PubMed Central PMCID: PMCPMC1571047.

21. Kataoka S, Yang R, Ishimaru Y, Matsunami H, Sevigny J, Kinnamon JC, et al. The candidate sour taste receptor, PKD2L1, is expressed by type III taste cells in the mouse. Chem Senses. 2008;33(3):243–54. doi: 10.1093/chemse/bjm083. PubMed PMID: 18156604; PubMed Central PMCID: PMCPMC2642677.

22. Oka Y, Butnaru M, von Buchholtz L, Ryba NJ, Zuker CS. High salt recruits aversive taste pathways. Nature. 2013;494(7438):472–5. Epub 2013/02/15. doi: 10.1038/nature11905. PubMed PMID: 23407495; PubMed Central PMCID: PMCPMC3587117.

23. Hacker K, Laskowski A, Feng L, Restrepo D, Medler K. Evidence for two populations of bitter responsive taste cells in mice. J Neurophysiol. 2008;99(3):1503–14. PubMed PMID: 18199819.

24. Clapp TR, Stone LM, Margolskee RF, Kinnamon SC. Immunocytochemical evidence for co-expression of Type III IP3 receptor with signaling components of bitter taste transduction. BMC Neurosci. 2001;2:6. PubMed PMID: 11346454.

25. Medler KF, Margolskee RF, Kinnamon SC. Electrophysiological characterization of voltage-gated currents in defined taste cell types of mice. J Neurosci. 2003;23(7):2608–17. PubMed PMID: 12684446.

26. Sukumaran SK, Lewandowski BC, Qin Y, Kotha R, Bachmanov AA, Margolskee RF. Whole transcriptome profiling of taste bud cells. Scientific reports. 2017;7(1):7595. doi: 10.1038/s41598-017-07746-z. PubMed PMID: 28790351; PubMed Central PMCID: PMCPMC5548921.

27. DeFazio RA, Dvoryanchikov G, Maruyama Y, Kim JW, Pereira E, Roper SD, et al. Separate populations of receptor cells and presynaptic cells in mouse taste buds. J Neurosci. 2006;26(15):3971–80. PubMed PMID: 16611813.

28. Richter TA, Caicedo A, Roper SD. Sour taste stimuli evoke Ca2+ and pH responses in mouse taste cells. J Physiol. 2003;547(Pt 2):475–83. PubMed PMID: 12562903.

29. Courtney E Wilson TF, Sue C Kinnamon Type III cells in anterior taste fields are more immunohistochemically diverse than those of posterior taste fields in mice. Chemical Senses. 2017;bjx055.

30. Tomchik SM, Berg S, Kim JW, Chaudhari N, Roper SD. Breadth of tuning and taste coding in mammalian taste buds. J Neurosci. 2007;27(40):10840–8. Epub 2007/10/05. doi: 27/40/10840 [pii] 10.1523/JNEUROSCI.1863-07.2007. PubMed PMID: 17913917.

31. Yoshida R, Miyauchi A, Yasuo T, Jyotaki M, Murata Y, Yasumatsu K, et al. Discrimination of taste qualities among mouse fungiform taste bud cells. J Physiol. 2009;587(Pt 18):4425–39. Epub 2009/07/23. doi: jphysiol.2009.175075 [pii] 10.1113/jphysiol.2009.175075. PubMed PMID: 19622604; PubMed Central PMCID: PMC2766648.

32. Shandilya J, Gao Y, Nayak TK, Roberts SG, Medler KF. AP1 transcription factors are required to maintain the peripheral taste system. Cell Death Dis. 2016;7(10):e2433. doi: 10.1038/cddis.2016.343. PubMed PMID: 27787515; PubMed Central PMCID: PMCPMC5133999.

33. Yang R, Crowley HH, Rock ME, Kinnamon JC. Taste cells with synapses in rat circumvallate papillae display SNAP-25-like immunoreactivity. J Comp Neurol. 2000;424(2):205–15. PubMed PMID: 10906698.

34. King CT, Garcea M, Spector AC. Glossopharyngeal nerve regeneration is essential for the complete recovery of quinine-stimulated oromotor rejection behaviors and central patterns of neuronal activity in the nucleus of the solitary tract in the rat. J Neurosci. 2000;20(22):8426–34. Epub 2000/11/09. PubMed PMID: 11069950.

35. King CT, Travers SP, Rowland NE, Garcea M, Spector AC. Glossopharyngeal nerve transection eliminates quinine-stimulated fos-like immunoreactivity in the nucleus of the solitary tract: implications for a functional topography of gustatory nerve input in rats. J Neurosci. 1999;19(8):3107–21. Epub 1999/04/07. PubMed PMID: 10191326.

36. Stratford JM, Finger TE. Central representation of postingestive chemosensory cues in mice that lack the ability to taste. J Neurosci. 2011;31(25):9101–10. Epub 2011/06/24. doi: 10.1523/JNEUROSCI.0404-11.2011. PubMed PMID: 21697361; PubMed Central PMCID: PMCPMC3131261.

37. Stratford JM, Thompson JA. MSG-Evoked c-Fos Activity in the Nucleus of the Solitary Tract Is Dependent upon Fluid Delivery and Stimulation Parameters. Chem Senses. 2016;41(3):211–20. Epub 2016/01/15. doi: 10.1093/chemse/bjv082. PubMed PMID: 26762887; PubMed Central PMCID: PMCPMC5006140.

38. Stratford JM, Thompson JA, Finger TE. Immunocytochemical organization and sour taste activation in the rostral nucleus of the solitary tract of mice. J Comp Neurol. 2017;525(2):271–90. Epub 2016/06/14. doi: 10.1002/cne.24059. PubMed PMID: 27292295; PubMed Central PMCID: PMCPMC5138149.

39. Hisatsune C, Yasumatsu K, Takahashi-Iwanaga H, Ogawa N, Kuroda Y, Yoshida R, et al. Abnormal Taste Perception in Mice Lacking the Type 3 Inositol 1,4,5-Trisphosphate Receptor. J Biol Chem. 2007;282(51):37225–31. PubMed PMID: 17925404.

40. Xu J, Lewandowski BC, Miyazawa T, Shoji Y, Yee K, Bryant BP. Spilanthol Enhances Sensitivity to Sodium in Mouse Taste Bud Cells. Chem Senses. 2019;44(2):91–103. Epub 2018/10/27. doi: 10.1093/chemse/bjy069. PubMed PMID: 30364996; PubMed Central PMCID: PMCPMC6350677.

41. Kimura K, Beidler LM. Microelectrode study of taste receptors of rat and hamster. J Cell Comp Physiol. 1961;58:131–9. Epub 1961/10/01. doi: 10.1002/jcp.1030580204. PubMed PMID: 14456037.

42. Sato T, Beidler LM. The response characteristics of rat taste cells to four basic taste stimuli. Comp Biochem Physiol A Comp Physiol. 1982;73(1):1–10. Epub 1982/01/01. doi: 10.1016/0300-9629(82)90083-4. PubMed PMID: 6127183.

43. Sato T, Beidler LM. Broad tuning of rat taste cells for four basic taste stimuli. Chem Senses. 1997;22(3):287–93. Epub 1997/06/01. doi: 10.1093/chemse/22.3.287. PubMed PMID: 9218141.

44. Tonosaki K, Funakoshi M. Intracellular taste cell responses of mouse. Comp Biochem Physiol A Comp Physiol. 1984;78(4):651–6. Epub 1984/01/01. doi: 10.1016/0300-9629(84)90611-x. PubMed PMID: 6149038.

45. Ozeki M. Conductance change associated with receptor potentials of gustatory cells in rat. J Gen Physiol. 1971;58(6):688–99. Epub 1971/12/01. doi: 10.1085/jgp.58.6.688. PubMed PMID: 5120394; PubMed Central PMCID: PMCPMC2226050.

46. Ozeki M, Sato M. Responses of gustatory cells in the tongue of rat to stimuli representing four taste qualities. Comp Biochem Physiol A Comp Physiol. 1972;41(2):391–407. Epub 1972/02/01. doi: 10.1016/0300-9629(72)90070-9. PubMed PMID: 4404316.

47. Caicedo A, Kim KN, Roper SD. Individual mouse taste cells respond to multiple chemical stimuli. J Physiol. 2002;544(Pt 2):501–9. PubMed PMID: 12381822.

48. Gilbertson TA, Boughter JD, Jr., Zhang H, Smith DV. Distribution of gustatory sensitivities in rat taste cells: whole-cell responses to apical chemical stimulation. J Neurosci. 2001;21(13):4931–41. PubMed PMID: 11425921.

49. Barretto RP, Gillis-Smith S, Chandrashekar J, Yarmolinsky DA, Schnitzer MJ, Ryba NJ, et al. The neural representation of taste quality at the periphery. Nature. 2015;517(7534):373–6. doi: 10.1038/nature13873. PubMed PMID: 25383521; PubMed Central PMCID: PMCPMC4297533.

50. Breza JM, Nikonov AA, Contreras RJ. Response latency to lingual taste stimulation distinguishes neuron types within the geniculate ganglion. J Neurophysiol. 2010;103(4):1771–84. doi: 10.1152/jn.00785.2009. PubMed PMID: 20107132; PubMed Central PMCID: PMCPMC2853290.

51. Frank ME, Lundy RF, Jr., Contreras RJ. Cracking taste codes by tapping into sensory neuron impulse traffic. Prog Neurobiol. 2008;86(3):245–63. doi: 10.1016/j.pneurobio.2008.09.003. PubMed PMID: 18824076; PubMed Central PMCID: PMCPMC2680288.

52. Hellekant G, Danilova V, Ninomiya Y. Primate sense of taste: behavioral and single chorda tympani and glossopharyngeal nerve fiber recordings in the rhesus monkey, Macaca mulatta. J Neurophysiol. 1997;77(2):978–93. doi: 10.1152/jn.1997.77.2.978. PubMed PMID: 9065862.

53. Wu A, Dvoryanchikov G, Pereira E, Chaudhari N, Roper SD. Breadth of tuning in taste afferent neurons varies with stimulus strength. Nat Commun. 2015;6:8171. doi: 10.1038/ncomms9171. PubMed PMID: 26373451; PubMed Central PMCID: PMCPMC4573454.

54. Zhang J, Jin H, Zhang W, Ding C, O’Keeffe S, Ye M, et al. Sour Sensing from the Tongue to the Brain. Cell. 2019;179(2):392–402 e15. Epub 2019/09/24. doi: 10.1016/j.cell.2019.08.031. PubMed PMID: 31543264.

55. Lemon CH, Smith DV. Neural representation of bitter taste in the nucleus of the solitary tract. J Neurophysiol. 2005;94(6):3719–29. doi: 10.1152/jn.00700.2005. PubMed PMID: 16107527.

56. Geran LC, Travers SP. Single neurons in the nucleus of the solitary tract respond selectively to bitter taste stimuli. J Neurophysiol. 2006;96(5):2513–27. PubMed PMID: 16899635.

57. Carleton A, Accolla R, Simon SA. Coding in the mammalian gustatory system. Trends Neurosci. 2010;33(7):326–34. doi: 10.1016/j.tins.2010.04.002. PubMed PMID: 20493563; PubMed Central PMCID: PMCPMC2902637.

58. Samuelsen CL, Gardner MP, Fontanini A. Thalamic contribution to cortical processing of taste and expectation. J Neurosci. 2013;33(5):1815–27. doi: 10.1523/JNEUROSCI.4026-12.2013. PubMed PMID: 23365221; PubMed Central PMCID: PMCPMC3711560.

59. Fontanini A, Katz DB. State-dependent modulation of time-varying gustatory responses. J Neurophysiol. 2006;96(6):3183–93. doi: 10.1152/jn.00804.2006. PubMed PMID: 16928791.

60. Katz DB, Simon SA, Nicolelis MA. Dynamic and multimodal responses of gustatory cortical neurons in awake rats. J Neurosci. 2001;21(12):4478–89. PubMed PMID: 11404435.

61. Accolla R, Bathellier B, Petersen CC, Carleton A. Differential spatial representation of taste modalities in the rat gustatory cortex. J Neurosci. 2007;27(6):1396–404. doi: 10.1523/JNEUROSCI.5188-06.2007. PubMed PMID: 17287514.

62. Zocchi D, Wennemuth G, Oka Y. The cellular mechanism for water detection in the mammalian taste system. Nat Neurosci. 2017;20(7):927–33. doi: 10.1038/nn.4575. PubMed PMID: 28553944.

63. Tu YH, Cooper AJ, Teng B, Chang RB, Artiga DJ, Turner HN, et al. An evolutionarily conserved gene family encodes proton-selective ion channels. Science. 2018;359(6379):1047–50. Epub 2018/01/27. doi: 10.1126/science.aao3264. PubMed PMID: 29371428; PubMed Central PMCID: PMCPMC5845439.

64. Teng B, Wilson CE, Tu YH, Joshi NR, Kinnamon SC, Liman ER. Cellular and Neural Responses to Sour Stimuli Require the Proton Channel Otop1. Curr Biol. 2019;29(21):3647–56 e5. Epub 2019/09/24. doi: 10.1016/j.cub.2019.08.077. PubMed PMID: 31543453.

65. Adler E, Hoon MA, Mueller KL, Chandrashekar J, Ryba NJ, Zuker CS. A novel family of mammalian taste receptors. Cell. 2000;100(6):693–702. PubMed PMID: 10761934.

66. Chandrashekar J, Hoon MA, Ryba NJ, Zuker CS. The receptors and cells for mammalian taste. Nature. 2006;444(7117):288–94. PubMed PMID: 17108952.

67. Chandrashekar J, Mueller KL, Hoon MA, Adler E, Feng L, Guo W, et al. T2Rs function as bitter taste receptors. Cell. 2000;100(6):703–11. PubMed PMID: 10761935.

68. Nelson G, Chandrashekar J, Hoon MA, Feng L, Zhao G, Ryba NJ, et al. An amino-acid taste receptor. Nature. 2002;416(6877):199–202. PubMed PMID: 11894099.

69. Nelson G, Hoon MA, Chandrashekar J, Zhang Y, Ryba NJ, Zuker CS. Mammalian sweet taste receptors. Cell. 2001;106(3):381–90. PubMed PMID: 11509186.

70. Stone LM, Barrows J, Finger TE, Kinnamon SC. Expression of T1Rs and gustducin in palatal taste buds of mice. Chem Senses. 2007;32(3):255–62. PubMed PMID: 17229761.

71. Yoshida R, Ninomiya Y. Taste information derived from T1R-expressing taste cells in mice. Biochem J. 2016;473(5):525–36. doi: 10.1042/BJ20151015. PubMed PMID: 26912569.

72. Hoon MA, Adler E, Lindemeier J, Battey JF, Ryba NJ, Zuker CS. Putative mammalian taste receptors: a class of taste-specific GPCRs with distinct topographic selectivity. Cell. 1999;96(4):541–51. PubMed PMID: 10052456.

73. Kim MR, Kusakabe Y, Miura H, Shindo Y, Ninomiya Y, Hino A. Regional expression patterns of taste receptors and gustducin in the mouse tongue. Biochem Biophys Res Commun. 2003;312(2):500–6. PubMed PMID: 14637165.

74. Kusakabe Y, Kim MR, Miura H, Shindo Y, Ninomiya Y, Hino A. Regional expression patterns of T1r family in the mouse tongue. Chem Senses. 2005;30 Suppl 1:i23–4. Epub 2005/03/02. doi: 10.1093/chemse/bjh094. PubMed PMID: 15738129.

75. Treesukosol Y, Blonde GD, Spector AC. T1R2 and T1R3 subunits are individually unnecessary for normal affective licking responses to Polycose: implications for saccharide taste receptors in mice. Am J Physiol Regul Integr Comp Physiol. 2009;296(4):R855–65. doi: 10.1152/ajpregu.90869.2008. PubMed PMID: 19158407; PubMed Central PMCID: PMC2698609.

76. Treesukosol Y, Smith KR, Spector AC. Behavioral evidence for a glucose polymer taste receptor that is independent of the T1R2+3 heterodimer in a mouse model. J Neurosci. 2011;31(38):13527–34. doi: 10.1523/JNEUROSCI.2179-11.2011. PubMed PMID: 21940444; PubMed Central PMCID: PMC3251913.

77. Treesukosol Y, Spector AC. Orosensory detection of sucrose, maltose, and glucose is severely impaired in mice lacking T1R2 or T1R3, but Polycose sensitivity remains relatively normal. Am J Physiol Regul Integr Comp Physiol. 2012;303(2):R218–35. doi: 10.1152/ajpregu.00089.2012. PubMed PMID: 22621968; PubMed Central PMCID: PMC3404635.

78. Zukerman S, Glendinning JI, Margolskee RF, Sclafani A. T1R3 taste receptor is critical for sucrose but not Polycose taste. Am J Physiol Regul Integr Comp Physiol. 2009;296(4):R866–76. doi: 10.1152/ajpregu.90870.2008. PubMed PMID: 19091911; PubMed Central PMCID: PMC2698610.

79. Kusuhara Y, Yoshida R, Ohkuri T, Yasumatsu K, Voigt A, Hubner S, et al. Taste responses in mice lacking taste receptor subunit T1R1. J Physiol. 2013;591(7):1967–85. doi: 10.1113/jphysiol.2012.236604. PubMed PMID: 23339178; PubMed Central PMCID: PMCPMC3624863.

80. Ohkuri T, Yasumatsu K, Horio N, Jyotaki M, Margolskee RF, Ninomiya Y. Multiple sweet receptors and transduction pathways revealed in knockout mice by temperature dependence and gurmarin sensitivity. Am J Physiol Regul Integr Comp Physiol. 2009;296(4):R960–71. doi: 10.1152/ajpregu.91018.2008. PubMed PMID: 19211717.

81. Chaudhari N, Landin AM, Roper SD. A metabotropic glutamate receptor variant functions as a taste receptor. Nat Neurosci. 2000;3(2):113–9. PubMed PMID: 10649565.

82. Chaudhari N, Maruyama Y, Roper S, Trubey K. Multiple pathways for signaling glutamate taste in rodents. Chem Senses. 2005;30 Suppl 1:i29–i30. PubMed PMID: 15738162.

83. Chaudhari N, Pereira E, Roper SD. Taste receptors for umami: the case for multiple receptors. Am J Clin Nutr. 2009;90(3):738S–42S. Epub 2009/07/03. doi: ajcn.2009.27462H [pii] 10.3945/ajcn.2009.27462H. PubMed PMID: 19571230.

84. Maruyama Y, Pereira E, Margolskee RF, Chaudhari N, Roper SD. Umami responses in mouse taste cells indicate more than one receptor. J Neurosci. 2006;26(8):2227–34. PubMed PMID: 16495449.

85. Delay ER, Hernandez NP, Bromley K, Margolskee RF. Sucrose and monosodium glutamate taste thresholds and discrimination ability of T1R3 knockout mice. Chem Senses. 2006;31(4):351–7. doi: 10.1093/chemse/bjj039. PubMed PMID: 16495435.

86. Eddy MC, Eschle BK, Delay ER. Comparison of the Tastes of L-Alanine and Monosodium Glutamate in C57BL/6J Wild Type and T1r3 Knockout Mice. Chem Senses. 2017;42(7):563–73. doi: 10.1093/chemse/bjx037. PubMed PMID: 28605507.

87. Pal Choudhuri S, Delay RJ, Delay ER. L-Amino Acids Elicit Diverse Response Patterns in Taste Sensory Cells: A Role for Multiple Receptors. PLoS One. 2015;10(6):e0130088. doi: 10.1371/journal.pone.0130088. PubMed PMID: 26110622; PubMed Central PMCID: PMCPMC4482487.

88. Pal Choudhuri S, Delay RJ, Delay ER. Metabotropic glutamate receptors are involved in the detection of IMP and L-amino acids by mouse taste sensory cells. Neuroscience. 2016;316:94–108. doi: 10.1016/j.neuroscience.2015.12.008. PubMed PMID: 26701297.

89. Shigemura N, Shirosaki S, Ohkuri T, Sanematsu K, Islam AA, Ogiwara Y, et al. Variation in umami perception and in candidate genes for the umami receptor in mice and humans. Am J Clin Nutr. 2009;90(3):764S–9S. Epub 2009/07/25. doi: ajcn.2009.27462M [pii] 10.3945/ajcn.2009.27462M. PubMed PMID: 19625681.

90. Sukumaran SK, Yee KK, Iwata S, Kotha R, Quezada-Calvillo R, Nichols BL, et al. Taste cell-expressed alpha-glucosidase enzymes contribute to gustatory responses to disaccharides. Proc Natl Acad Sci U S A. 2016;113(21):6035–40. Epub 2016/05/11. doi: 10.1073/pnas.1520843113. PubMed PMID: 27162343; PubMed Central PMCID: PMCPMC4889361.

91. Dvoryanchikov G, Hernandez D, Roebber JK, Hill DL, Roper SD, Chaudhari N. Transcriptomes and neurotransmitter profiles of classes of gustatory and somatosensory neurons in the geniculate ganglion. Nat Commun. 2017;8(1):760. Epub 2017/10/04. doi: 10.1038/s41467-017-01095-1. PubMed PMID: 28970527; PubMed Central PMCID: PMCPMC5624912.

92. Carr CE, Konishi M. A circuit for detection of interaural time differences in the brain stem of the barn owl. J Neurosci. 1990;10(10):3227–46. Epub 1990/10/01. PubMed PMID: 2213141.

93. Joris PX, Smith PH, Yin TC. Coincidence detection in the auditory system: 50 years after Jeffress. Neuron. 1998;21(6):1235–8. Epub 1999/01/12. PubMed PMID: 9883717.

94. Oertel D, Bal R, Gardner SM, Smith PH, Joris PX. Detection of synchrony in the activity of auditory nerve fibers by octopus cells of the mammalian cochlear nucleus. Proc Natl Acad Sci U S A. 2000;97(22):11773–9. Epub 2000/10/26. doi: 10.1073/pnas.97.22.11773. PubMed PMID: 11050208; PubMed Central PMCID: PMCPMC34348.

95. Ala-Laurila P, Rieke F. Coincidence detection of single-photon responses in the inner retina at the sensitivity limit of vision. Curr Biol. 2014;24(24):2888–98. Epub 2014/12/03. doi: 10.1016/j.cub.2014.10.028. PubMed PMID: 25454583; PubMed Central PMCID: PMCPMC4269560.

96. Sakmann B. From single cells and single columns to cortical networks: dendritic excitability, coincidence detection and synaptic transmission in brain slices and brains. Exp Physiol. 2017;102(5):489–521. Epub 2017/02/01. doi: 10.1113/EP085776. PubMed PMID: 28139019; PubMed Central PMCID: PMCPMC5435930.

97. Brill MF, Meyer A, Rossler W. It takes two-coincidence coding within the dual olfactory pathway of the honeybee. Front Physiol. 2015;6:208. Epub 2015/08/19. doi: 10.3389/fphys.2015.00208. PubMed PMID: 26283968; PubMed Central PMCID: PMCPMC4516877.

98. Gupta N, Singh SS, Stopfer M. Oscillatory integration windows in neurons. Nat Commun. 2016;7:13808. Epub 2016/12/16. doi: 10.1038/ncomms13808. PubMed PMID: 27976720; PubMed Central PMCID: PMCPMC5171764.

99. Wang L, Gillis-Smith S, Peng Y, Zhang J, Chen X, Salzman CD, et al. The coding of valence and identity in the mammalian taste system. Nature. 2018;558(7708):127–31. Epub 2018/06/01. doi: 10.1038/s41586-018-0165-4. PubMed PMID: 29849148; PubMed Central PMCID: PMCPMC6201270.

100. Pfaff DW. Theoretical consideration of cross-fiber pattern coding in the neural signalling of pheromones and other chemical stimuli. Psychoneuroendocrinology. 1975;1:79–93.

101. Hegg CC, Jia C, Chick WS, Restrepo D, Hansen A. Microvillous cells expressing IP3 receptor type 3 in the olfactory epithelium of mice. Eur J Neurosci. 2010;32(10):1632–45. doi: 10.1111/j.1460-9568.2010.07449.x. PubMed PMID: 20958798; PubMed Central PMCID: PMCPMC4331646.

102. Han SK, Mancino V, Simon MI. Phospholipase Cbeta 3 mediates the scratching response activated by the histamine H1 receptor on C-fiber nociceptive neurons. Neuron. 2006;52(4):691–703. doi: 10.1016/j.neuron.2006.09.036. PubMed PMID: 17114052.

103. Xie W, Samoriski GM, McLaughlin JP, Romoser VA, Smrcka A, Hinkle PM, et al. Genetic alteration of phospholipase C beta3 expression modulates behavioral and cellular responses to mu opioids. Proc Natl Acad Sci U S A. 1999;96(18):10385–90. PubMed PMID: 10468617; PubMed Central PMCID: PMCPMC17897.

104. Hacker K, Medler KF. Mitochondrial calcium buffering contributes to the maintenance of Basal calcium levels in mouse taste cells. J Neurophysiol. 2008;100(4):2177–91. doi: 10.1152/jn.90534.2008. PubMed PMID: 18684902; PubMed Central PMCID: PMCPMC2576209.

105. Laskowski AI, Medler KF. Sodium-calcium exchangers contribute to the regulation of cytosolic calcium levels in mouse taste cells. J Physiol. 2009;587(Pt 16):4077–89. doi: 10.1113/jphysiol.2009.173567. PubMed PMID: 19581381; PubMed Central PMCID: PMCPMC2756439.

106. Maliphol AB, Garth DJ, Medler KF. Diet-induced obesity reduces the responsiveness of the peripheral taste receptor cells. PLoS One. 2013;8(11):e79403. doi: 10.1371/journal.pone.0079403. PubMed PMID: 24236129; PubMed Central PMCID: PMC3827352.

107. Rebello MR, Maliphol AB, Medler KF. Ryanodine Receptors Selectively Interact with L Type Calcium Channels in Mouse Taste Cells. PLoS One. 2013;8(6):e68174. doi: 10.1371/journal.pone.0068174. PubMed PMID: 23826376; PubMed Central PMCID: PMC3694925.

108. Rebello MR, Medler KF. Ryanodine receptors selectively contribute to the formation of taste-evoked calcium signals in mouse taste cells. Eur J Neurosci. 2010;32(11):1825–35. Epub 2010/10/20. doi: 10.1111/j.1460-9568.2010.07463.x. PubMed PMID: 20955474; PubMed Central PMCID: PMC2994989.

109. Szebenyi SA, Laskowski AI, Medler KF. Sodium/calcium exchangers selectively regulate calcium signaling in mouse taste receptor cells. J Neurophysiol. 2010;104(1):529–38. Epub 2010/05/14. doi: jn.00118.2010 [pii] 10.1152/jn.00118.2010. PubMed PMID: 20463203; PubMed Central PMCID: PMC2904227.

110. Bartel DL, Sullivan SL, Lavoie EG, Sevigny J, Finger TE. Nucleoside triphosphate diphosphohydrolase-2 is the ecto-ATPase of type I cells in taste buds. J Comp Neurol. 2006;497(1):1–12. doi: 10.1002/cne.20954. PubMed PMID: 16680780; PubMed Central PMCID: PMCPMC2212711.

111. Harrer MI, Travers SP. Topographic organization of Fos-like immunoreactivity in the rostral nucleus of the solitary tract evoked by gustatory stimulation with sucrose and quinine. Brain Res. 1996;711(1-2):125–37. Epub 1996/03/04. PubMed PMID: 8680855.

112. Martin LE, Nikonova LV, Kay K, Paedae AB, Contreras RJ, Torregrossa AM. Salivary proteins alter taste-guided behaviors and taste nerve signaling in rat. Physiol Behav. 2017;184:150–61. doi: 10.1016/j.physbeh.2017.11.021. PubMed PMID: 29162505.

113. Torregrossa AM, Nikonova L, Bales MB, Villalobos Leal M, Smith JC, Contreras RJ, et al. Induction of salivary proteins modifies measures of both orosensory and postingestive feedback during exposure to a tannic acid diet. PLoS One. 2014;9(8):e105232. doi: 10.1371/journal.pone.0105232. PubMed PMID: 25162297; PubMed Central PMCID: PMCPMC4146545.

114. Preacher KJ. Calculation for the chi-square test: An interactive calculation tool for chi-square tests of goodness of fit and independence [Computer software]. 2001. Available from http://quantpsy.org.].

